# ISRIB blunts the integrated stress response by allosterically antagonising the inhibitory effect of phosphorylated eIF2 on eIF2B

**DOI:** 10.1101/2020.09.28.316653

**Authors:** Alisa F. Zyryanova, Kazuhiro Kashiwagi, Claudia Rato, Heather P. Harding, Ana Crespillo-Casado, Luke A. Perera, Ayako Sakamoto, Madoka Nishimoto, Mayumi Yonemochi, Mikako Shirouzu, Takuhiro Ito, David Ron

## Abstract

The small molecule ISRIB antagonises the activation of the integrated stress response (ISR) by phosphorylated translation initiation factor 2, eIF2(αP). ISRIB and eIF2(αP) bind distinct sites in their common target, eIF2B, a guanine nucleotide exchange factor (GEF) for eIF2. We have found that ISRIB-mediated acceleration of eIF2B activity in vitro is observed preferentially in the presence of eIF2(αP) and is attenuated by mutations that desensitise eIF2B to the inhibitory effects of eIF2(αP). ISRIB’s efficacy as an ISR inhibitor in cells also depends on presence of eIF2(αP). Cryo-EM showed that engagement of both eIF2B regulatory sites by two eIF2(αP) molecules remodels both the ISRIB-binding pocket and the pockets that would engage eIF2α during active nucleotide exchange, thereby discouraging both binding events. In vitro, eIF2(αP) and ISRIB reciprocally opposed each other’s binding to eIF2B. These findings point to antagonistic allostery in ISRIB action on eIF2B, culminating in inhibition of the ISR.

## Introduction

Under diverse stressful conditions, the α subunit of eukaryotic translation initiation factor 2 (eIF2) is phosphorylated on serine 51. This converts eIF2, the substrate of eIF2B, to an inhibitor of eIF2B; a guanine nucleotide exchange factor (GEF) that normally reactivates the eIF2 heterotrimer by accelerating the release of GDP from the eIF2γ subunit and its exchange with GTP (Ranu and London, 1979; de Haro et al., 1996). By depleting ternary complexes of eIF2, GTP and initiator methionyl-tRNA (Met-tRNA_i_) in the cell, eIF2α phosphorylation attenuates the translation of most mRNAs, with important effects on protein synthesis. On the other hand, translation of a small number of mRNAs is increased in an eIF2 phosphorylation-dependent manner. As the latter encode potent transcription factors, the production of phosphorylated eIF2 [eIF2(αP)] is coupled with a conserved gene expression programme referred to as the integrated stress response (ISR) (Harding et al., 2003).

The ISR is a homeostatic pathway that contributes to organismal fitness (Pakos-Zebrucka et al., 2016). However, in some circumstances its heightened activity is associated with unfavourable outcomes, motivating a search for ISR inhibitors. When applied to cells or administered to animals, the druglike small molecule, ISRIB, disrupts the ISR (Sidrauski et al., 2013), and has been reported to exert beneficial effects in models of neurodegeneration (Halliday et al., 2015; Zhu et al., 2019), head injury (Chou et al., 2017) and dysmyelination (Wong et al., 2018; Abbink et al., 2019).

ISRIB does not affect the levels of eIF2(αP), indicating a site of action downstream of this common effector. ISRIB-resistant mutations were mapped genetically to the eIF2B β and δ subunits and disrupt the high affinity binding of ISRIB (*K_d_* ~10 nM) to a pocket on the surface of eIF2B (Sekine et al., 2015; Tsai et al., 2018; Zyryanova et al., 2018), demonstrating that eIF2B is ISRIB’s target.

eIF2B is a large (~500 kDa) decamer, assembled from two sets of five subunits (Kashiwagi et al., 2016). It has two catalytic sites, each comprised of the bipartite eIF2Bε subunit whose two lobes embrace the nucleotide-binding eIF2γ subunit enforcing a conformation that favours GDP dissociation and exchange with GTP. Engagement of eIF2 in this catalytically productive conformation depends on binding of the N-terminal lobe of the eIF2α subunit in a pocket between the β and δ subunits of eIF2B, ~100 Å from the catalytic site (Kashiwagi et al., 2019; Kenner et al., 2019). When eIF2 is phosphorylated, the N-terminal lobe of P-eIF2α engages eIF2B at an alternative site, between eIF2Bα and eIF2Bδ (Adomavicius et al., 2019; Gordiyenko et al., 2019; Kashiwagi et al., 2019; Kenner et al., 2019): a catalytically non-productive binding mode that inhibits eIF2B’s nucleotide exchange activity (*in cis* or *trans*, Kashiwagi et al., 2019). ISRIB binds a single and distinct site in eIF2B, at its centre of symmetry, the interface between the β and δ subunits of eIF2B (Tsai et al., 2018; Zyryanova et al., 2018).

It stands to reason that ISRIB inhibits the ISR by promoting the nucleotide exchange activity of eIF2B. Indeed, when added to crude preparations of eIF2B, ISRIB accelerates exchange of GDP nucleotide on its substrate eIF2 (Sekine et al., 2015; Sidrauski et al., 2015). A simple mechanism has been proposed to account for such stimulation: decameric eIF2B consists of one α_2_ dimer and two βδγε tetramers; by binding across the interface between the two tetramers, ISRIB favours eIF2B decamer assembly and stability. According to this model, which is well supported by a study of eIF2B assembly in vitro, ISRIB inhibits the ISR by increasing the effective concentration of active, decameric eIF2B (Tsai et al., 2018).

Accelerated assembly of eIF2B as ISRIB’s mode of action would be favoured by the presence of a large pool of unassembled eIF2B subunits in the cell. Yet, fractionation of mammalian cell lysates by density gradient centrifugation has not suggested the existence of large pools of precursor complexes of eIF2B subunits (Sidrauski et al., 2015; Zyryanova et al., 2018). Being a slow fractionation method, density gradient centrifugation might fail to detect a pool of precursors migrating at their predicted position in the gradient, if the precursors were in a rapid equilibrium with the assembled decamers. However, the finding that ISRIB has little to no effect on the nucleotide exchange activity of pure eIF2B decamers, in vitro (Tsai et al., 2018), speaks against ISRIB increasing active enzyme concentration by stabilising the decamer in such a rapid equilibrium regime.

These considerations prompted us to examine the evidence for alternative modes of ISRIB action. Given the physical distance between the ISRIB binding site and the sites by which eIF2B engages the various components of eIF2, such action would necessarily proceed by allosteric mechanisms. Here we report on biochemical, structural and cell-based findings that ISRIB allosterically antagonises the inhibitory effect of eIF2(αP) on eIF2B guanine nucleotide exchange activity to inhibit the ISR.

## Results

### ISRIB accelerates eIF2B guanine nucleotide exchange activity in presence of eIF2(αP)

To assess the effects of ISRIB on the eIF2B guanine nucleotide exchange activity in isolation of phosphorylated eIF2, we loaded BODIPY-GDP onto eIF2(α^S51A^) isolated from FreeStyle 293-F cells expressing mutated eIF2 that harbours the S51A mutation in the α subunit. We then utilised eIF2(α^S51A^) as a substrate in a fluorescence-based nucleotide exchange assay with recombinant human eIF2B; the latter purified from bacteria. As reported previously (Tsai et al., 2018), ISRIB only minimally accelerated the exchange of nucleotide mediated by eIF2B in an assay devoid of eIF2(αP) (Figure 1A). Introduction of eIF2(αP) into the assay attenuated the nucleotide exchange activity directed towards the non-phosphorylatable eIF2(α^S51A^)•BODIPY-GDP. This effect was significantly, though only partially, reversed by ISRIB (Figure 1B), as observed previously (Wong et al., 2018).

**Figure 1.**
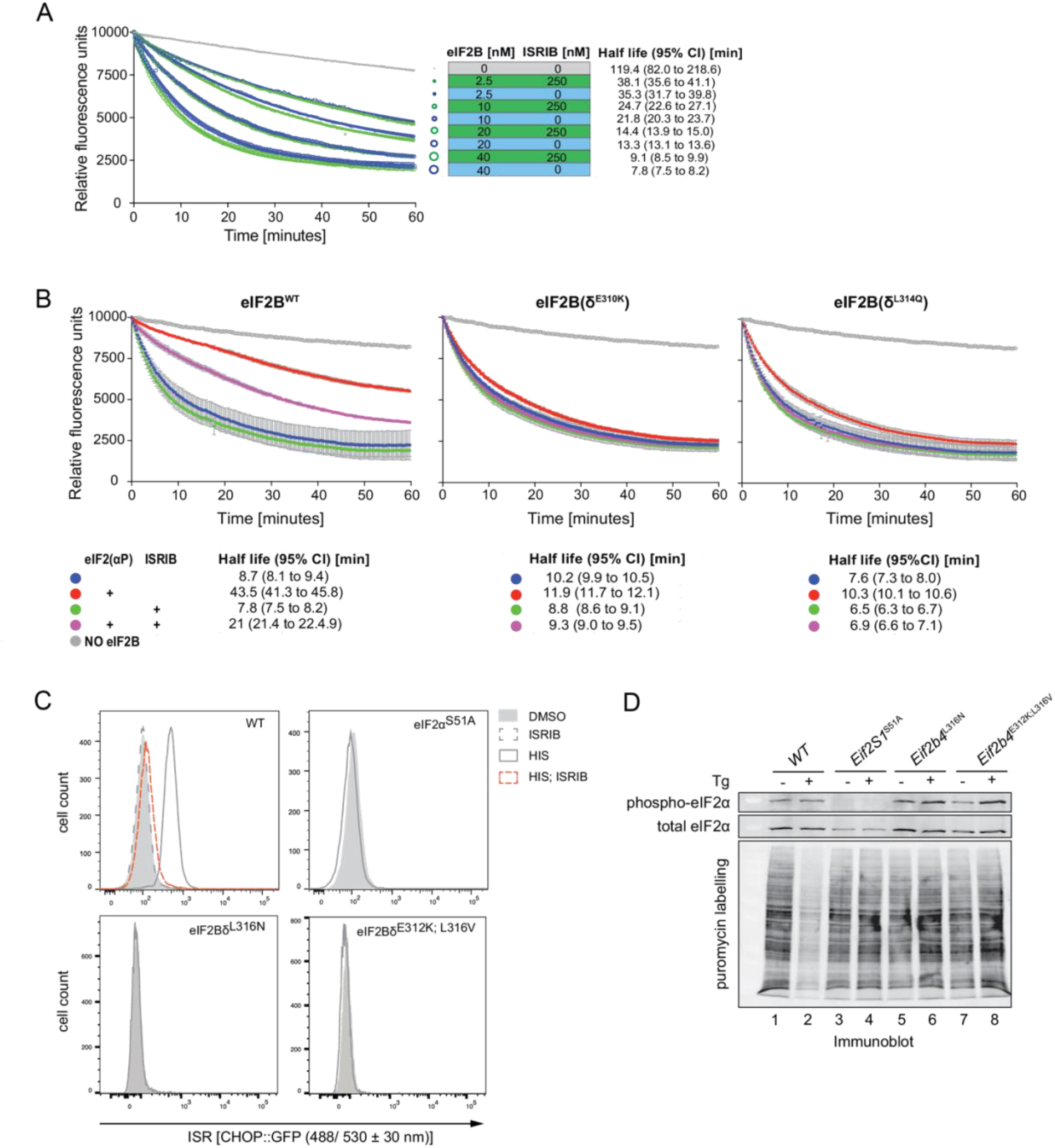
ISRIB accelerates eIF2B guanine nucleotide exchange activity selectively in presence of eIF2(αP). A) Plots of eIF2B guanine nucleotide exchange activity as reflected in time-dependent decrease in fluorescence of BODIPY-FL-GDP bound to non-phosphorylatable eIF2(α^S51A^) (125 nM). Blue traces lack and green traces include ISRIB (250 nM), the grey trace lacks eIF2B. The size of the symbol reflects the concentration of eIF2B in the assay. All the data points of a representative experiment performed in duplicate are shown. The half-life of GDP binding with 95% confidence interval (CI) for each plot is indicated (observation reproduced three times). B) As in “A” but utilising a fixed concentration of wildtype or the indicated ISR-insensitive eIF2B mutants (40 nM), BODIPY-FL-GDP bound to non-phosphorylatable eIF2(α^S51A^) (125 nM) and where indicated, phosphorylated eIF2 (1 μM) and ISRIB (250 nM). Plotted are the mean fluorescence values ± SD of an experiment performed in technical triplicate. C) In vivo characterisation of the ISR in wildtype and mutant CHO cells of the indicated genotype. Shown are histograms of the activity of CHOP::GFP (an ISR reporter gene) in untreated cells and cells in which the eIF2α kinase GCN2 had been activated by L-histidinol in the absence or presence of ISRIB. Shown is a representative experiment reproduced three times. D) Estimates of protein synthesis rates in CHO cells of the indicated genotype before and after activation of the eIF2α kinase PERK by thapsigargin. The lower panel is an anti-puromycin immunoblot of whole cell lysates, in which the intensity of the puromycinylated protein signal reports on rates of protein synthesis. The upper panels are immunoblots of phosphorylated eIF2α and of total eIF2α.

The *trans*-inhibitory effect of eIF2(αP) observed in vitro was markedly attenuated by mutations in human eIF2Bδ residues, E310K and L314Q (Figure 1B), as reported previously (Kimball et al., 1998). When introduced into the genome of cultured CHO cells (by CRISPR/Cas9-mediated homologous recombination), mutations in the corresponding hamster residues (eIF2Bδ E312 and L316) imparted an ISR-attenuated phenotype, as reflected in the blunted stress-induced activation of the ISR-responsive CHOP::GFP reporter gene (Figure 1C) and in the blunted repression of protein synthesis, normally observed in stressed cells (Figure 1D). These findings are consistent with the phenotype of corresponding mutations in yeast, GCD2^E377K^ and GCD2^L381Q^ (Pavitt et al., 1997). In vitro, equivalent substitutions in human eIF2Bδ, E310K and L314Q, also blunted the response to ISRIB that was observed with wildtype eIF2B in presence of eIF2(αP) (Figure 1B). Together these observations indicate that, in vitro, ISRIB reverses an inhibitory effect of eIF2(αP) on the guanine nucleotide exchange activity of eIF2B that is relevant to the activation of the ISR in vivo. Notably, these observations are difficult to reconcile with a mechanism of action that relies solely on ISRIB-mediated stabilisation of the eIF2B decamer. If such stabilisation were a significant contributor, it would effectively increase the enzyme concentration, and register as a proportional increase in the nucleotide exchange activity even in the absence of eIF2(αP).

### eIF2(αP) induces an eIF2B conformation inimical to ISRIB binding

To better understand the basis for ISRIB’s ability to antagonise the *trans*-inhibitory effect of eIF2(αP) observed in the experiments above, we solved the cryo-electron microscopy (cryoEM) structures of human apo eIF2B and in complex with eIF2(αP) trimer (Figure 2, Supplement 1 and Table S1). Though the overall binding modes of eIF2(αP) in these structures are similar to previous ones (Adomavicius et al., 2019; Gordiyenko et al., 2019;

Kashiwagi et al., 2019; Kenner et al., 2019), the new structures highlight variation among the complexes, namely between complexes with one eIF2(αP) trimer, in which only the α subunit is resolved (termed the α^P^1 complex), complexes with two eIF2(αP) trimers, in which two molecules of the α subunit are resolved at both sides (the α^P^2 complex), and complexes with two eIF2(αP) trimers, in which both the α and γ subunits are resolved at one side and only the α subunit resolved at the other side (the α^P^γ complex). The subunits conformations in the α^P^γ complex are almost identical to those in previous structures (Kashiwagi et al., 2019), but resolution is improved.

When the apo eIF2B and α^P^γ complex structures are compared, the binding of eIF2(αP), which is contributed by E310 and L314 residues of eIF2Bδ, induced a noticeable movement of the βδγε tetrameric unit of eIF2B (Figure 2A). This rotation widens the pocket between eIF2Bβ and eIF2Bδ that would otherwise accommodate the N-terminal domain of eIF2α in the catalytically productive conformation (Figure 2B and figure 2 supplement 2) (Kashiwagi et al., 2019; Kenner et al., 2019). A widened gap between eIF2Bβ and eIF2Bδ appears to be a conserved feature, as the conformation of yeast eIF2B, bound by two eIF2(αP) trimers more closely resembles the human α^P^γ complex than human apo eIF2B (Gordiyenko et al, 2019).

**Figure 2.**
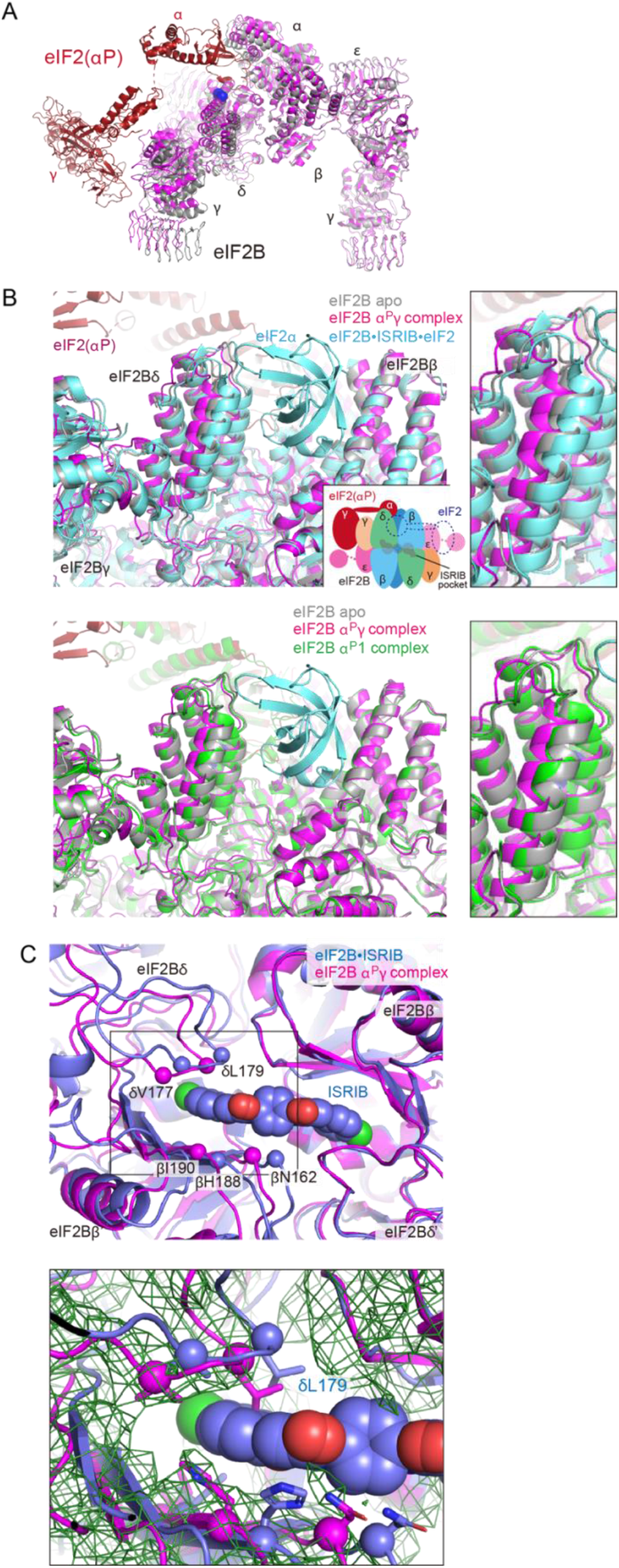
eIF2(αP) and ISRIB associate with different conformations of eIF2B. A) Overlay of the eIF2B apo structure (grey) and eIF2B in complex with two eIF2(αP) trimers (the α^P^γ complex; eIF2B in pink and eIF2(αP) in red). The blue spheres show the position of eIF2Bδ^E310^ and δ^L314^. Structures are aligned by their eIF2Bβ subunits. B) Different effects on the interface that accommodates the N-terminal domain of eIF2α in the catalytically productive conformation in complexes containing one (α^P^1) or two (α^P^γ) bound eIF2(αP) trimers. Upper panel: an overlay of the productive eIF2B•ISRIB•eIF2 complex (cyan, PDB: 6O85), eIF2B apo structure (grey) and the α^P^γ complex (magenta) aligned by their eIF2Bβ subunits. For clarity, only the N-terminal lobe of eIF2α of the eIF2B•ISRIB•eIF2 complex, is shown. A cartoon of the complex (inset) is provided for orientation. Lower panel: similar alignment of the apo structure, α^P^γ structure and α^P^1 structure, in green. The N-terminal lobe of eIF2α in the eIF2B•ISRIB•eIF2 complex is shown for reference (cyan). C) Deformation of the ISRIB binding pocket in eIF2B with two bound eIF2(αP) trimers. Upper panel: The eIF2B•ISRIB complex structure (PDB: 6CAJ) is shown in blue and α^P^γ complex in magenta aligned by their eIF2Bδ’ subunits. Key residues lining the pocket, including eIF2B δ^L179^ and β^H188^ that are known to affect the binding or action of ISRIB, are highlighted as spheres and sticks. Lower panel: The EM density map for the α^P^γ complex is shown in dark green.

Because the accommodation of the N-terminal domain of unphosphorylated eIF2α induces closure at this interface (PDB: 6O85) (Kenner et al, 2019) (Figure 2B and figure 2, supplement 2), the binding of eIF2(αP) is seen to antagonise catalytically productive binding. A trend in the same direction was also observed in a previous structure of eIF2B complexed with isolated phosphorylated eIF2α subunit (P-eIF2α) (PDB: 6O9Z) (Kenner et al., 2019), but the displacement of the βδγε tetrameric unit is more pronounced in our presently-determined α^P^γ complex structure (Figure 2, supplement 2). Therefore, the γ subunit of eIF2(αP) seems to contribute to this structural rearrangement of eIF2B’s subunits. The contribution of contacts between the γ subunit of eIF2(αP) and eIF2Bγ, noted here in the α^P^γ complex, to these transitions remains unclear as some rotation in the same direction is already noted in the α^P^2 complex (Figure 2, supplement 2B).

The rearrangement of eIF2B induced by eIF2(αP) also affects the pocket for ISRIB formed by two sets of eIF2Bβ and eIF2Bδ subunits (Tsai et al., 2018; Zyryanova et al., 2018). Especially, the side-chains of eIF2Bδ^L179^ and eIF2Bβ^H188^, key residues involved in ISRIB action and binding (Sekine et al., 2015; Tsai et al., 2016; Zyryanova et al., 2018), are shifted compared with the eIF2BoISRIB complex (PDB: 6CAJ) (Figure 2C and figure 2, supplement 2C). This is predicted to compromise the accommodation of ISRIB’s symmetrically-positioned aryl groups. In contrast, the structural change of this pocket induced by the binding of unphosphorylated eIF2 is marginal and compatible with co-binding of ISRIB (Figure 2, supplement 2A) (Kenner et al, 2019).

Compared to the α^P^γ and α^P^2 complexes that contain two eIF2(αP) trimers and induce movement in eIF2B as described above, the rearrangement observed in the α^P^1 complex, which contains only one eIF2(αP) trimer, is subtler. Although the eIF2Bα subunit is marginally displaced following the accommodation of eIF2(αP) (Figure 2, supplement 2D), there is negligible shifting in the other parts of eIF2B, including the interface for unphosphorylated eIF2 and the pocket for ISRIB (Figures 2B&C, and figure 2, supplement 2B&2C). Therefore, the aforementioned movement induced by eIF2(αP) was accentuated by accommodation of the second eIF2(αP) trimer. The coupling between eIF2(αP) binding at its regulatory sites and the progressive deformation of the ISRIB binding pocket brought about by sequential binding of two eIF2(αP) trimers sets the stage for a competition, whereby ISRIB-mediated stabilisation of its pocket is propagated in a reciprocal manner to the eIF2(αP) binding sites. ISRIB is expected to be especially antagonistic towards engagement of a second eIF2(αP) trimer, hence discouraging eIF2B from assuming its most inhibited conformation.

### Antagonism between eIF2(αP) and ISRIB binding to eIF2B in vitro

These structural insights predict mutually antagonistic binding of eIF2(αP) and ISRIB to eIF2B. To test this prediction, we deployed a fluorescence polarisation-based assay to measure the binding of a FAM-labelled ISRIB to eIF2B in vitro (Zyryanova et al., 2018). The binding of the small FAM-ISRIB (MW ~ 1 kDa) to the much larger wildtype or ISR-insensitive mutants eIF2B(δ^E310K^) and eIF2B(δ^L314Q^) (MW ~ 500 kDa) results in a similar marked increase in the fluorescence polarisation signal (Figure 3A).

**Figure 3.**
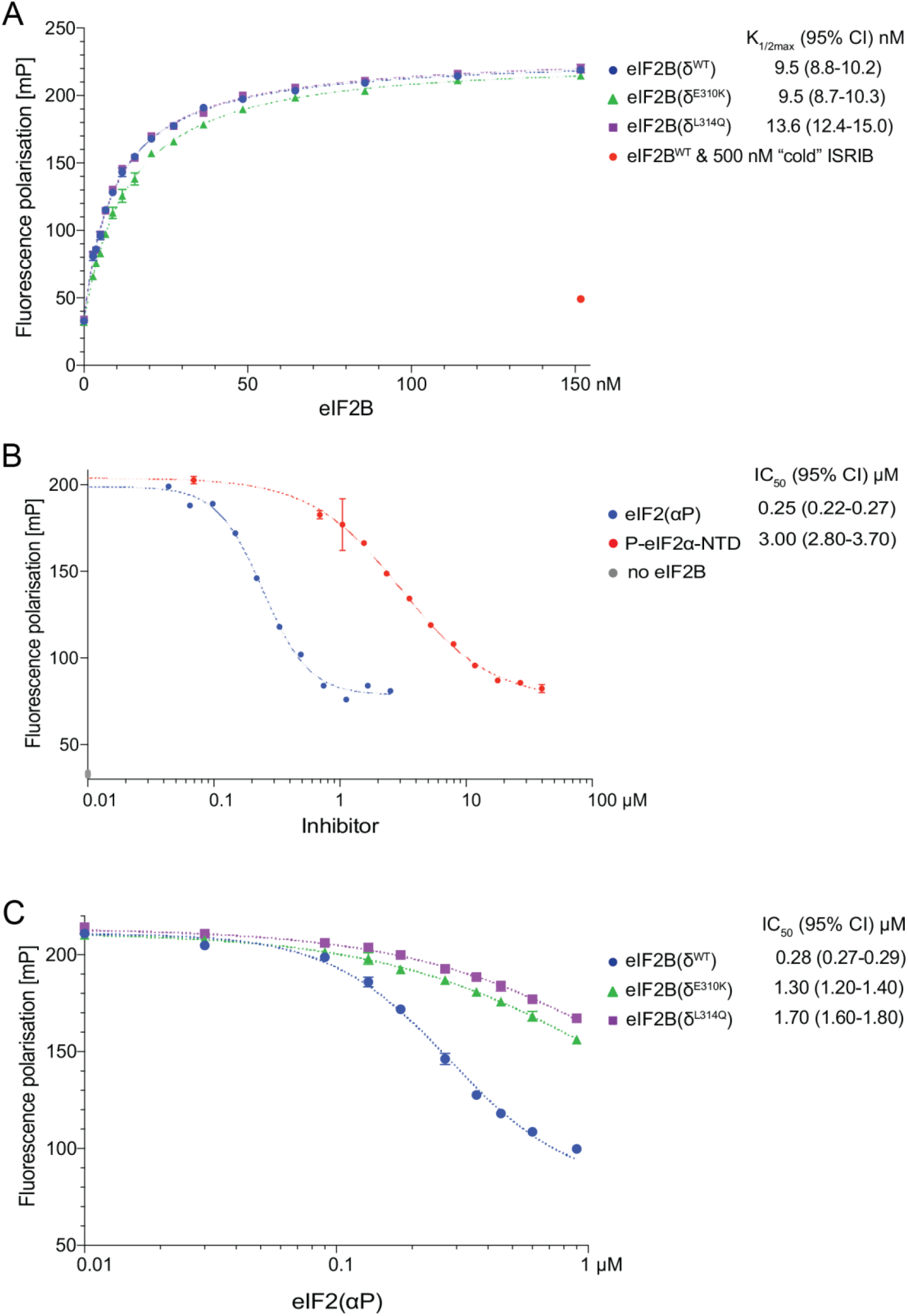
Phosphorylated eIF2 attenuates FAM-ISRIB binding to eIF2B. A) Plot of fluorescence polarisation signals (mean ± SD, n=3) arising from samples of FAM conjugated ISRIB (2.5 nM) incubated with varying concentrations of wildtype or mutant eIF2B. Where indicated, 500 nM unlabelled ISRIB was added as a competitor. *K_1/2max_* with 95% confidence intervals (CI) is shown. B) Plot of fluorescence polarisation signals, at equilibrium, arising from FAM-ISRIB bound to wildtype eIF2B in presence of the indicated concentration of the N-terminal lobe of phosphorylated eIF2α [P-eIF2α, mean ± SD (n = 3)] or eIF2(αP) trimer. The data were fitted by non-linear regression analysis to a “log[inhibitor] vs. response four parameter” model and the IC_50_ values with 95% CI are shown. C) As in “B” above, plot of fluorescence polarisation signals, at equilibrium, arising from FAM-ISRIB bound to wildtype or mutant eIF2B (100 nM) in presence of the indicated concentration of eIF2(αP) trimer [mean ± SD (n = 3)]. IC_50_ values with 95% CI are shown.

Challenge of the eIF2BoFAM-ISRIB complex with eIF2(αP) resulted in a concentration dependent decrease in the fluorescence polarisation signal at steady state with an IC_50_ ~0.25 μM (Figure 3B and 3C). The Hill slope of the reaction 2.4, suggests a cooperative process, consistent with the requirement for binding of two molecules of eIF2(αP) for the displacement of FAM-ISRIB. The isolated N-terminal lobe of the phosphorylated eIF2α subunit (P-eIF2α-NTD) also displaced FAM-ISRIB from eIF2B, but with an IC_50_ that was >10-fold higher (3.0 μM, Figure 3B). Complexes formed between FAM-ISRIB and the ISR-insensitive eIF2B(δ^E310K^) and eIF2B(δ^L314Q^) were more resistant to the inhibitory effect of eIF2(αP) (Figure 3C) despite wildtype affinity for FAM-ISRIB (Figure 3A).

The eIF2BoFAM-ISRIB complex is maintained dynamically: unlabelled ISRIB displaced FAM-ISRIB from eIF2B with *k_off_* of 0.74 min^-1^ (green diamond trace in Figure 4A). The presence of eIF2 did not affect the stability of the eIF2B•ISRIB complex over time (open blue circle trace in Figure 4B top). However, introduction of the PERK kinase into the assay (in presence of ATP), which resulted in the gradual phosphorylation of eIF2, led to a time-dependent loss of the fluorescence polarisation signal (blue diamond trace in Figure 4B top). The PERKdependent decline in signal was enzyme concentration-dependent, correlated with eIF2 phosphorylation (compare blue, lilac and red square traces in Figure 4, supplement 1A) and recovered in a time dependent manner by introducing phosphatases that dephosphorylated eIF2 (lilac and orange square traces in Figure 4, supplement 1B). These features attest to the dynamism and reversible nature of this in vitro representation of the ISR in the presence of ISRIB.

**Figure 4.**
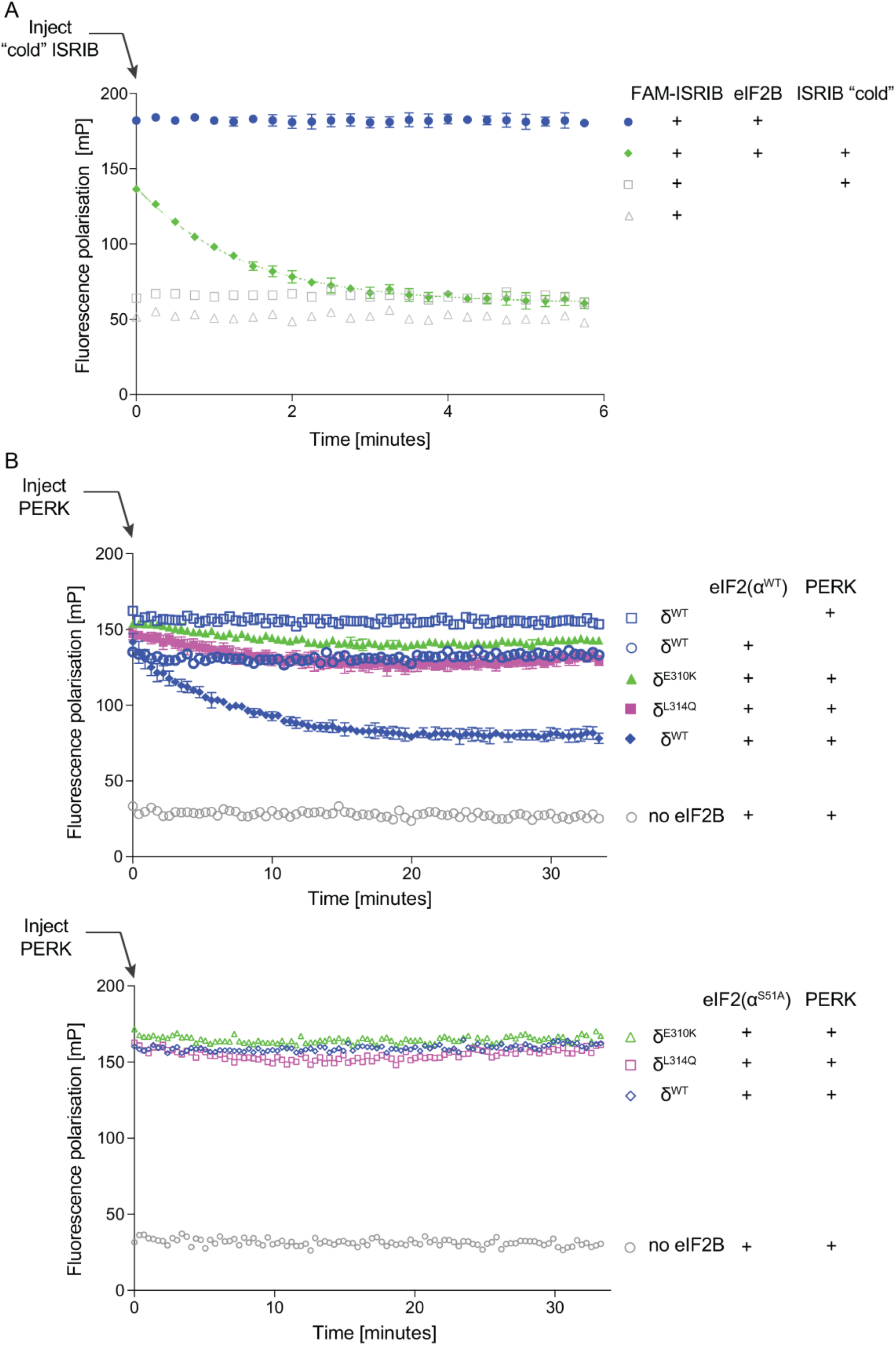
Phosphorylated eIF2 attenuates FAM-ISRIB binding to eIF2B on a timescale consistent with the ISR. A) Plot of the time-dependent change in fluorescence polarisation of FAM-ISRIB bound to wildtype eIF2B, following injection of 1 μM unlabelled ISRIB at t = 0 [green diamonds, mean ± SD, n = 3) and the fit of the first six minutes to a first order decay reaction (k_*off*_ = 0.74 min^-1^, 95% CI 0.68 - 0.9 min^-1^, R^2^ 0.9823), dotted green line]. Control samples, unchallenged by cold ISRIB (blue circles, mean ± SD, n = 3) and reference samples (n = 1) from the same experiment are shown. B) Plot of time-dependent change in fluorescence polarisation of FAM-ISRIB bound to wildtype or ISR-insensitive mutant eIF2Bs (δ^E310K^ or δ^L314Q^) (60 nM) in presence or absence of 600 nM unphosphorylated eIF2 (wildtype, top panel) or non-phosphorylatable eIF2(α^S51A^) (mutant, bottom panel). Where indicated, at t = 0 the eIF2α kinase PERK was introduced to promote a pool of eIF2(αP). Shown are the mean ± SD (n = 3) of the fluorescence polarisation values of the PERK-injected samples. The traces were fitted to a first order decay reaction. [WT: *t*_1/2_ 4.7 minutes (95% CI, 4.5 - 6.7, R^2^ 0.9344); δ^E310K^: *t*_1/2_ 153 minutes and δ^L314Q^: *t*_1/2_ 92 minutes (both with a poor fit to first order decay, R^2^ < 0.5)].

The time-dependent PERK-mediated loss of fluorescence polarisation signal was not evident when wildtype eIF2 was replaced by a mutant eIF2(α^S51A^) that is unable to serve as a substrate for PERK (Figure 4B bottom). eIF2 phosphorylation-mediated loss of fluorescence polarisation signal arising from FAM-ISRIB binding to wildtype eIF2B was attenuated by the ISR-insensitive mutants, eIF2Bδ^E310K^ and eIF2Bδ^L314Q^ (Figure 4B top, green triangles and lilac squares, respectively). This last finding implies that the eIF2(αP)-mediated rearrangement of eIF2B that lowers its affinity for ISRIB is dependent on the same contacts that are required for eIF2(αP)’s ability to trigger the ISR.

To assess the impact of ISRIB on the association of phosphorylated eIF2 with eIF2B, we turned to biolayer interferometry (BLI). The biotinylated phosphorylated N-terminal lobe of eIF2α (P-eIF2α), immobilised via streptavidin to a BLI sensor, gave rise to a greater optical signal when reacted with fully assembled eIF2B decamers in solution compared to either eIF2B^βδγε^ tetramers (Tsai et al., 2018) (Figure 5A), or the ISR-insensitive mutants, eIF2B(δ^E310K^ or δ^L314Q^) (Figure 5B). These features suggested that physiologically-relevant contacts between eIF2B and P-eIF2α contributed significantly to the BLI signal.

**Figure 5.**
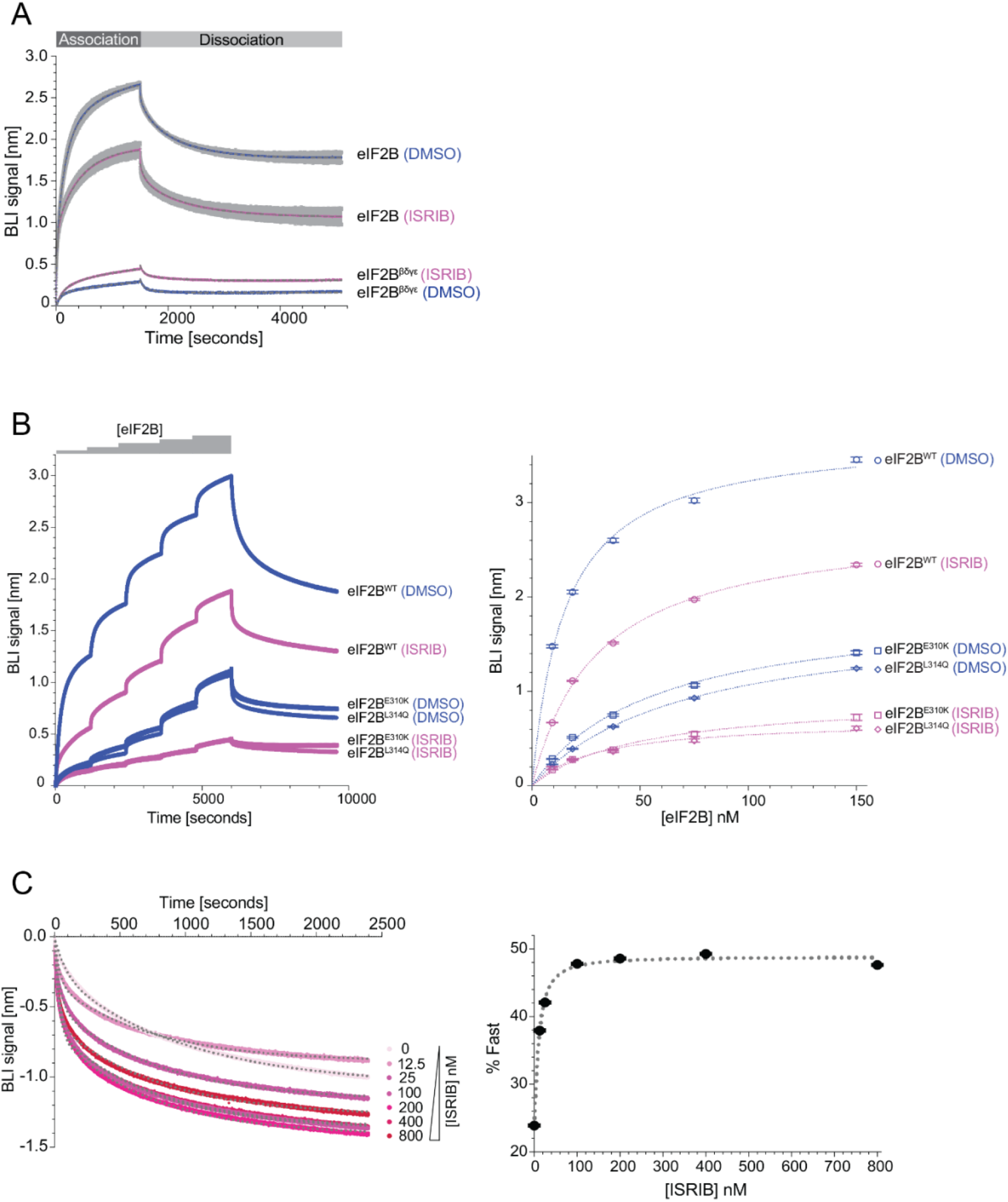
ISRIB inhibits binding of eIF2B to the N-terminal lobe of phosphorylated eIF2α. A) Biolayer interferometry (BLI) traces of the association and dissociation phases of eIF2B decamers (100 nM) or eIF2B^βδγε^ tetramers (400 nM) in the absence (DMSO, in blue) or presence of ISRIB (1 μM, in pink) to and from biotinylated P-eIF2α (N-terminal lobe) immobilised on the BLI probe. The fits to a 2-phase association and a 2-phase dissociation model are indicated by the grey dashed line. Shown are mean ± SD (n = 3) of the sample of eIF2B decamers with and without ISRIB and all the data points of the tetramer samples from a representative experiment conducted three times. B) Left: BLI traces of consecutive association phases and a terminal dissociation phase of wildtype and the indicated eIF2B mutants in the absence (DMSO, in blue) or presence of ISRIB (1 μM, in pink) as in “A”. The probe was reacted with escalating concentrations of eIF2B (9 to 150 nM) ± ISRIB, before dissociation in the respective buffer. Right: Plots of the mean and 95% confidence interval of the plateau values of the association phases (obtained by fitting the data from the traces on the left to a 2-phase association non-linear regression model) against the concentration of eIF2B. The dotted line reports on the fit of plots to a specific binding with Hill slope = 1 non-linear regression model. Shown are all the data points of a representative experiment performed three times. C) Left: The time-dependent change in BLI signal in the dissociation phase of eIF2B (previously-associated in absence of ISRIB) from biotinylated P-eIF2α in the presence of escalating concentration of ISRIB. The mean (and 95% confidence intervals) of the fraction of the dissociation attributed to the fast phase (%Fast) was calculated by fitting the dissociation traces to a biphasic model (the fit is indicated by the grey dotted lines overlaying the data points in the traces). Shown are traces from a representative experiment performed three times. Right: Plot of the %Fast of the dissociation reactions to the left, as a function of ISRIB concentration. The plot was fitted to an [Agonist] vs. response - variable slope (four parameters) non-linear regression model (dotted line) yielding an EC_50_ of 10.5 (95% CI, 4.3 - 16.3) nM.

The presence of ISRIB attenuated the association of P-eIF2α with eIF2B, across a broad range of eIF2B concentrations. The association and dissociation reactions detected by BLI were multi-phasic and therefore likely comprised of more than one binding event. Nonetheless, both the association phase and the dissociation phase gave a very good fit to double exponential models. This enabled estimation of ISRIB’s effect on both eIF2B’s steady state binding *[K1/2* max of eIF2B-dependent BLI signal increased from 15.2 nM in the absence of ISRIB to 29.8 nM in its presence (Figure 5B)] and on the kinetics of eIF2B dissociation [ISRIB increased the PercentFast dissociation from 23.9% to 49.3%, with an EC_50_ of 10.5 nM, (Figure 5C)].

Together, these experiments point to antagonism between engagement of eIF2(αP) and ISRIB as eIF2B ligands, at their respective distinct sites. Given that ISRIB binding to eIF2B favours, whilst eIF2(αP) binding disfavours, binding of eIF2 as a substrate for the nucleotide exchange, these findings suggest a plausible mechanism whereby ISRIB-mediated stabilisation of the active conformation of the eIF2B decamer allosterically antagonises the ISR.

### Attenuated ISRIB action in cells lacking eIF2(αP)

To learn more about the relative roles of allostery and eIF2B assembly in ISRIB’s action in vivo, we turned to cells lacking phosphorylated eIF2. The ISR in CHO cells in which the wildtype eIF2α encoding gene had been replaced by an *Eif2S1^S51A^* mutant allele is unresponsive to manipulations that activate eIF2α kinases (Crespillo-Casado et al., 2017, and Figure 1C). However, in these *Eif2S1^S51A^* mutant cells, CRISPR/Cas9 disruption of *Eif2b* subunit-encoding genes activated the ISR, as reflected in the time dependent emergence of a population of cells expressing high levels of CHOP::GFP (Figure 6A). Despite the progressive loss of viability attendant upon depletion of eIF2B (reflected in the decline in the CHOP::GFP bright subpopulation, observed 96 hours after transduction with gene-specific guides and Cas9), this assay enabled the measurement of ISRIB’s effect on the ISR in absence of any phosphorylated eIF2.

**Figure 6.**
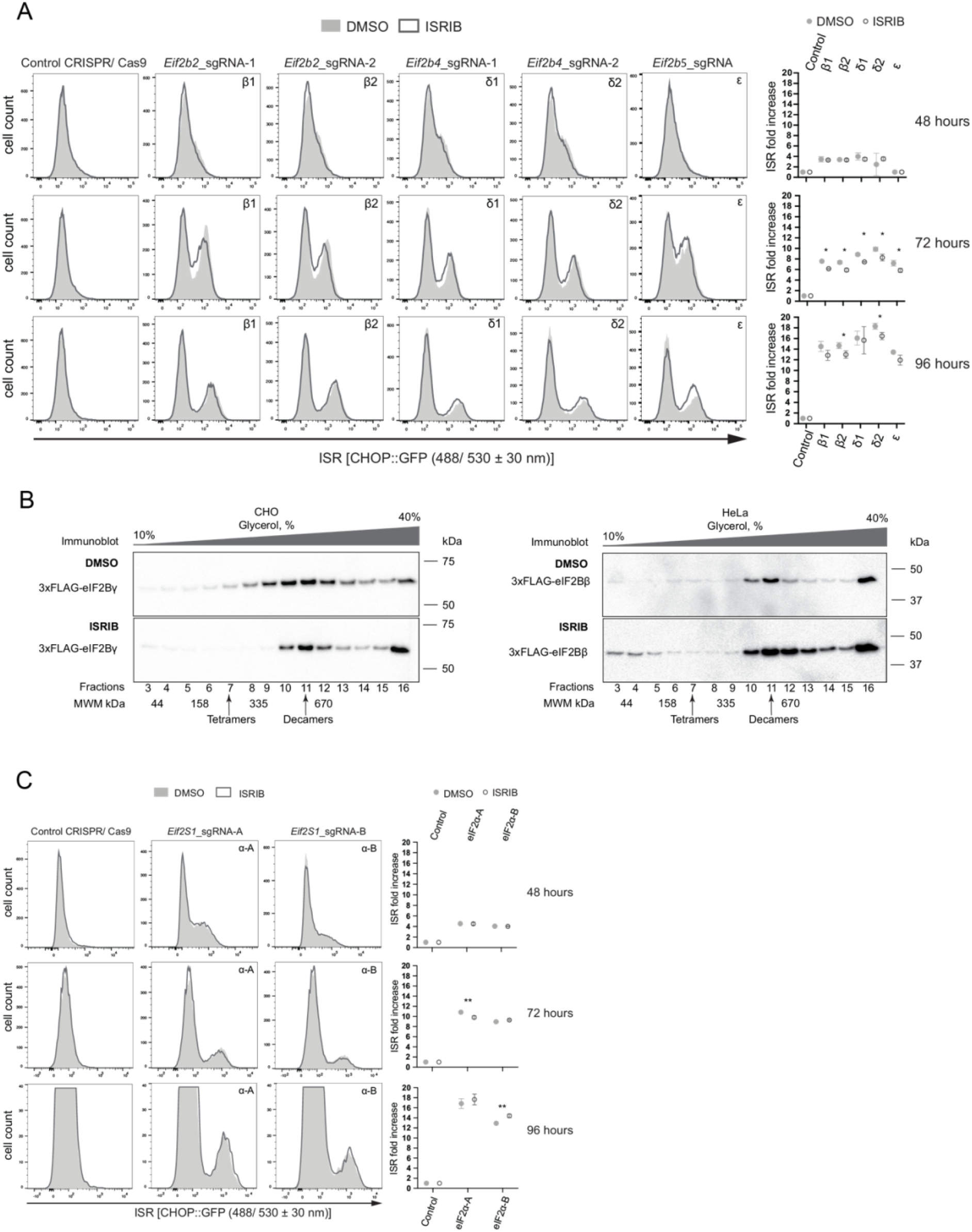
Attenuated ISRIB action in cells lacking eIF2(αP). A) In vivo characterisation of the ISR in *Eif2S1^S51A^* mutant CHO cells (lacking phosphorylatable eIF2α) depleted of eIF2B subunits by CRISPR/Cas9 targeting of their encoding genes. Two different guides for either beta (β1, β2) or delta (δ1, δ2) subunit, and one guide for epsilon subunit (ε) were transfected separately. Where indicated the cells were exposed continuously to ISRIB (1 μM), commencing at the point of transduction with the CRISPR/Cas9 encoding plasmids and continued until harvest. Shown are histograms of the CHOP::GFP ISR reporter in populations of cells 48, 72 & 96 hours following eIF2B gene targeting (± ISRIB) from a representative experiment performed three times. The mean ± SD of the ratio of fluorescent signal of the ISR induced population (the right peak on the histograms) to the non-induced (left peak) from three independent experiments are plotted (*P < 0.05 Student’s t-test). B) Immunoblot of 3 x FLAG-tagged endogenous eIF2Bγ detected with anti-FLAG M2 antibodies in CHO or 3 x FLAG-tagged endogenous eIF2Bβ in HeLa cell lysates that were either treated with DMSO (top panel) or ISRIB (bottom panel) and resolved on a 10 - 40% glycerol density gradient in a buffer of physiological salt concentration. The position of reference proteins of the indicated molecular weight in this gradient is indicated below the image and the arrows point to the predicted position of eIF2B(βδγε) tetramers and eIF2B(α)_2_(βδγε)_2_ decamers. C) As in “A” above, but following CRISPR/Cas9-mediated depletion of eIF2B’s substrate eIF2, by targeting the *Eif2S1* gene encoding its α subunit. Two different guides, eIF2α-A and eIF2α-B, were transfected separately. Shown is a representative experiment performed twice (**P < 0.05 Student’s t-test).

ISRIB had only a very modest (albeit statistically significant) inhibitory effect on the magnitude of the ISR induced by eIF2B subunit depletion, despite comparable levels of CHOP::GFP activation to those observed in L-histidinol-treated wildtype cells (Figure 6A, compare to Figure 1C). This finding - a weak ISRIB effect under conditions of eIF2B subunit depletion and no eIF2 phosphorylation - is consistent with the in vitro observation that in the absence of eIF2(αP) ISRIB only weakly stimulated the nucleotide exchange activity of eIF2B, even when the enzyme’s concentration was lowered by dilution (Figure 1A and Tsai et al., 2018). Thus, it appears that whilst ISRIB’s high affinity binding to eIF2B can undoubtedly stabilise both the assembled decamer and its intermediates in vitro (Tsai et al., 2018), the contribution of this mechanism to its action in CHO cells is rather limited.

The aforementioned considerations are in keeping with the finding that density gradients of cell lysates do not support the existence of two substantial pools of eIF2B subunits: one of assembled decamers and another of unassembled eIF2B intermediates (Sidrauski et al., 2015; Zyryanova et al., 2018). Nonetheless, scrutiny of immunoblots of density gradients of both CHO and HeLa cell lysates (prepared under physiological salt conditions) does suggest a small but conspicuous pool of tagged endogenous eIF2Bγ (or eIF2Bβ) subunits migrating in the gradient at the position expected of an eIF2Bβδγε tetramer (MW 229 kDa). In both cell types, this minor pool of putative assembly intermediates appears to be depleted by ISRIB (Figure 6B). The latter observation is consistent with a measure of ISRIB-mediated acceleration of eIF2B assembly and also suggests that the limited pool of unassembled intermediates may account for ISRIB’s limited residual effect on the ISR, observed in *Eif2S1^S51A^* mutant cells.

Depletion of eIF2B subunits, by interfering with their production (through CRISPR/Cas9-mediated gene disruption), is predicted to cut off the supply of even this modest pool of precursors and deprive ISRIB of an opportunity to increase eIF2B’s activity via enhanced assembly. Therefore, to gage the importance of accelerated assembly to ISRIB action in a different experimental system, we depleted cells of eIF2B’s substrate by inactivating the *Eif2S1* gene (in the *EifS1^S51A^* cells). As expected, this manipulation also activated the ISR, despite the absence of any phosphorylated eIF2α. However, in this scenario too the stimulatory effect of ISRIB was very modest (compare Figure 6C with Figure 1C). Together these findings suggest that in CHO cells ISRIB reversal of the ISR is realised mostly through its ability to antagonise the effects of eIF2(αP) on pre-existing eIF2B decamers, via the allosteric mechanism described here.

## Discussion

Comparing experimental systems containing and lacking phosphorylated eIF2 demonstrated the importance of eIF2(αP) to unveil ISRIB’s ability to promote nucleotide exchange in vitro or ISR inhibition in cells. This correlates with structural observations whereby ISRIB binding is associated with a conformation of eIF2B conducive to binding of eIF2 as a substrate, whilst eIF2(αP) binding is associated with a different conformation of eIF2B with an altered ISRIB binding pocket. Binding of ISRIB and eIF2(αP) to eIF2B appears mutually antagonistic. The inhibitory effect of eIF2(αP) on ISRIB binding to eIF2B is played out in vitro over a time frame consistent with the dynamics of the ISR (Harding et al., 2000). Reciprocally, a robust and rapid inhibitory effect of ISRIB on the association between the N-terminal lobe of phosphorylated eIF2α and eIF2B is also noted. Together, these findings point to an allosteric component of ISRIB action, whereby its binding to eIF2B stabilises the latter in a conformation that is relatively resistant to eIF2(αP). Given eIF2(αP)’s role as the major known upstream inducer of the ISR, this proposed allosteric mechanism goes some way to explaining ISRIB’s ability to antagonise this cellular response to stress.

Dependence of ISRIB-mediated stimulation of eIF2B’s nucleotide exchange activity on the presence of eIF2(αP) in vitro (Wong et al., 2018), and an apparent incompatibility between ISRIB binding to eIF2B and the conformation imposed on eIF2B by eIF2(αP) (Gordiyenko et al., 2019) had both been suggested previously. Furthermore, whilst particles of ternary complexes of eIF2B•ISRIB•eIF2 (PDB: 6O85) are readily attainable (Kenner et al., 2019), efforts to assemble similar particles with eIF2B, ISRIB and eIF2(αP) have been unsuccessful. Our findings here unify these earlier clues, supporting the conclusion that eIF2(αP) and ISRIB are incompatible ligands of eIF2B.

Structural analysis suggests at least two components to the aforementioned incompatibility. The first relates to changes imposed on the regulatory cleft of eIF2B by the binding of the N-terminal domain of phosphorylated eIF2α between the eIF2Bα and δ subunits. Our structural observations suggest that such changes are enforced cooperatively by the binding of two molecules of the N-terminal domain of phosphorylated eIF2α at both symmetrical regulatory sites of eIF2B (Figure 7A). The importance of these contacts, to toggle eIF2B to a non-ISRIB binding mode, is demonstrated by an attenuated effect of eIF2(αP) on the binding of FAM-ISRIB to eIF2B in the context of the ISR-desensitising mutants, eIF2B(δ^E310K^) and eIF2B(δ^L314Q^). The second relates to a role for the βγ lobe of eIF2(αP), since the rearrangement of eIF2B was more prominent in structures containing the phosphorylated eIF2 trimer compared with those of eIF2B complexed with isolated P-eIF2α. This finding is mirrored in the ~10-fold lower IC_50_ of the eIF2(αP) trimer, compared with the isolated N-terminal domain of phosphorylated α subunit, in the inhibition of FAM-ISRIB binding to eIF2B. It is tempting to speculate that contacts between the γ subunit of eIF2(αP) and eIF2Bγ observed in some classes of particles in the cryo-EM images may stabilise the inhibited eIF2BoeIF2(αP) complex, but this issue has yet to be examined experimentally.

**Figure 7.**
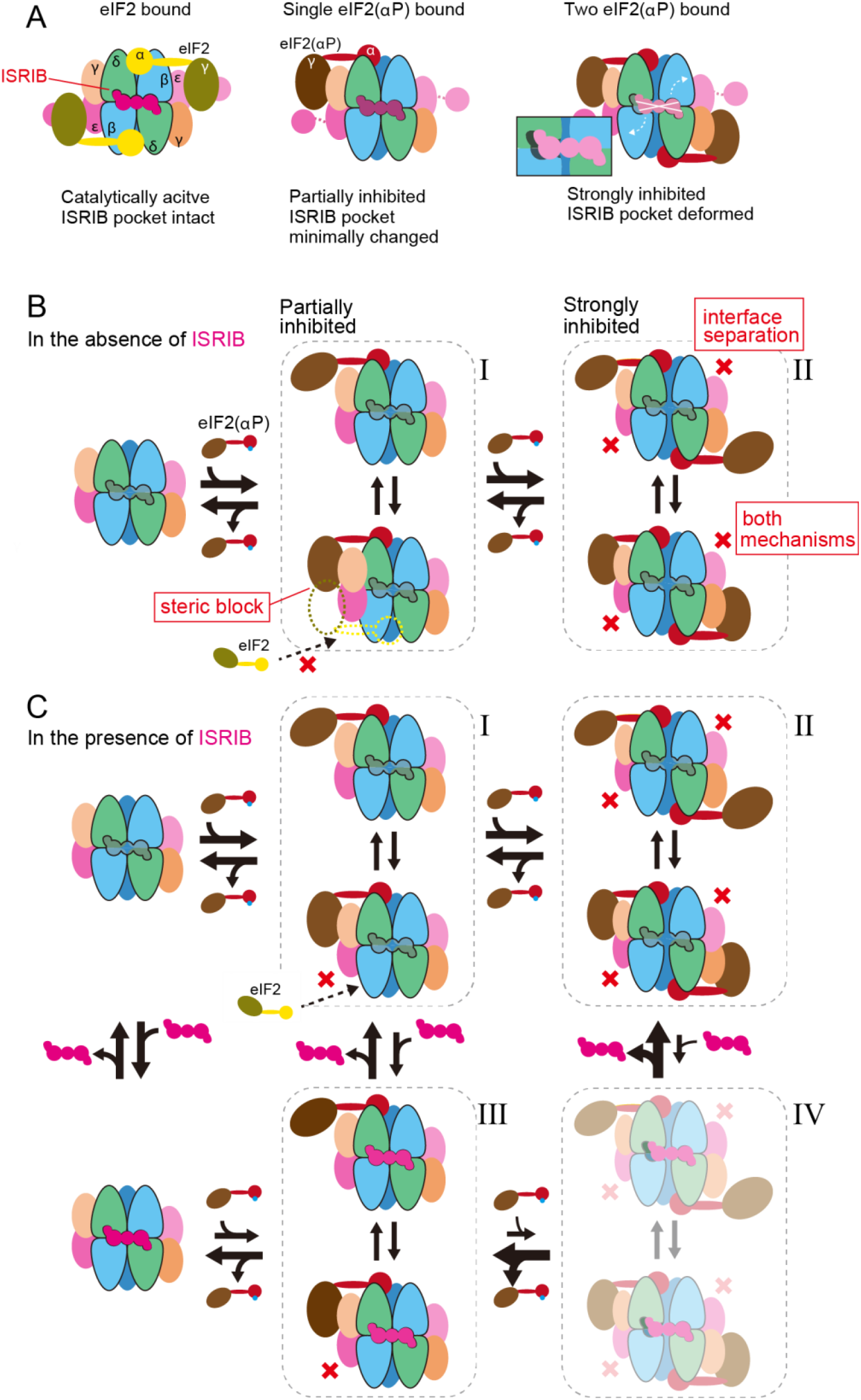
A Model of the functional consequences of the antagonism between eIF2(αP) and ISRIB binding to eIF2B. A) Cartoon of the ISRIB binding pocket in the active (ground state) of eIF2B (left), eIF2B bound by one eIF2(αP) trimer (centre) and eIF2B bound by two eIF2(αP) (right) trimers. B) The binding of one eIF2(αP) trimer partially inhibits the catalytic activity *in trans* by a **steric block** of the active site induced by the docking of the γ subunit of eIF2(αP) onto eIF2Bγ (state I). Binding of a second eIF2(αP) trimer exposes the second active site of eIF2B to a similar steric block but also interferes with catalytic activity *in cis* through **interface separation** that deforms both pockets for productive binding of eIF2α, strongly inhibiting the catalytic activity of eIF2B (state II). C) The presence of ISRIB hierarchically antagonises the binding of eIF2(αP). Weaker antagonism towards one bound eIF2(αP) trimer (states I & III), as ISRIB’s binding pocket is less deformed under these circumstances. However, the stronger deformation of the ISRIB binding pocket that is coupled to the binding of a second eIF2(αP) trimer sets up a competition whereby state IV is most strongly disfavoured by ISRIB. By stabilising the ground state and the partially inhibited state III, ISRIB pulls the equilibrium away from the strongly inhibited state II, thus antagonising the ISR. High enough concentrations of eIF2(αP) competitively over-ride this inhibition, consistent with observations that in cells ISRIB is a partial antagonist of the ISR.

It was previously noted in the eIF2BoeIF2(αP) complex that the phosphorylated eIF2’s βγ lobe blocks access of an unphosphorylated eIF2 trimer (bound *in trans)* to eIF2B’s catalytic site (Kashiwagi et al., 2019). This mechanism favours partial inhibition of the catalytic activity of eIF2B when it accommodates a single eIF2(αP) trimer (state I in Figure 7B). In this study we note that the movement of eIF2Bδ away from eIF2Bβ, observed in the eIF2Bo[eIF2(αP)]2 complex with two bound eIF2(αP) trimers (the α^P^2 and α^P^γ structures), widens the groove that could otherwise productively engage the N-terminal domain of unphosphorylated eIF2α subunit of a third eIF2 trimer as a substrate, and is thus predicted to destabilise an active enzyme-substrate complex also *in cis*. It seems reasonable to imagine that eIF2B assumes this strongly inhibited state in presence of higher concentrations of eIF2(αP) (state II in Figure 7B).

Simultaneous binding of eIF2(αP) to both regulatory sites deforms the ISRIB binding pocket. It is plausible that the rigid, largely helical structure of eIF2B’s regulatory domain [comprised of the helical extensions of its α, β and δ subunits (Kuhle et al., 2015)] contributes to this allosteric coupling, rendering the concurrent binding of ISRIB with two molecules of eIF2(αP) unlikely. At low levels of eIF2(αP) the competition thus set up enables ISRIB to antagonise the transition of eIF2B from the fully active to the strongly inhibited state (the one most incompatible with ISRIB binding, Figure 7C) thereby dampening the cellular response to increasing levels of eIF2 phosphorylation. The kinetic parameters governing this antagonistic allostery have yet to be determined. However, the observation that ISRIB is only a partial antagonist of the ISR (Halliday et al., 2015; Rabouw et al., 2019) suggests that at high enough concentrations eIF2(αP) can outcompete ISRIB.

To parse the contribution of the allosteric antagonism between ISRIB and eIF2(αP) demonstrated here from the role of ISRIB in accelerating assembly of eIF2B decamers (Tsai et al., 2018), we experimentally activated the ISR in *Eif2S1^S51A^* cells lacking P-eIF2α by transient genetic manipulations that deplete their pool of ternary eIF2oGTPoMet-tRNA_i_ complexes. In absence of eIF2(αP), the limited velocity of the nucleotide exchange reaction, imposed by either substrate or enzyme depletion, resulted in an ISR that was only weakly antagonised by ISRIB. These findings argue against an important role for accelerated assembly or eIF2B stabilisation in ISRIB action in CHO cells. This conclusion also fits with the paucity of evidence for a substantial pool of eIF2B precursors for ISRIB to draw on and accelerate assembly of eIF2B in cells under basal conditions. Nor is there evidence to suggest that the eIF2B decamer is in a rapid exchange equilibrium containing a significant fraction of eIF2Bβδγε tetramers and eIF2Bα dimers, as such an equilibrium would be expected to be skewed by ISRIB towards the active GEF decamer in vitro and in vivo, even in absence of eIF2(αP). It is noteworthy that we have not ruled out the possibility that eIF2(αP) may itself perturb the decamer-tetramer equilibrium of eIF2B, thus uncovering the potential stabilising activity of ISRIB as a basis for ISR inhibition. However, efforts to otherwise favour the dissolution of decamers in the absence of eIF2(αP), by dilution to a concentration 20-100 fold lower than that found in cells (53-293 nM, Hein et al., 2015) (Figure 1A) or by in vivo depletion of individual eIF2B subunits (Figure 6A), did not result in the manifestation of marked ISRIB effects.

The relative contribution of allostery and stabilisation to ISRIB’s action may be different in cells with mutations in eIF2B that lower its enzymatic activity by destabilising the decamer. Stabilisation may therefore contribute importantly to the salubrious role of ISRIB (and the related compound 2BAct) in cellular and animal models of the myelinopathy associated with eIF2B mutations (Wong et al., 2018; Abbink et al., 2019; Wong et al., 2019). It is also possible that stabilisation of the eIF2B decamer may be important in other contexts, such as regulation of eIF2B by subunit phosphorylation (Wang et al., 2001) or in other cell types, which may have significant unassembled pools of eIF2B precursors (Hodgson et al., 2019)

Given the discovery of ISRIB’s role as an allosteric regulator of eIF2B presented here, it is interesting to contemplate the potential contribution of decamer assembly/stability and allostery to the action of other ligands of the ISRIB pocket - be they yet-to-be discovered physiological regulators of translation or drugs. Particularly interesting is the question of whether ligands of the ISRIB pocket can be discovered that stabilise the inactive conformation of eIF2B - the one imposed on it by eIF2(αP). If ISRIB attains its effects in cells largely by allostery, acting on a cellular pool of stable eIF2B decamers (as our findings here suggest), such anti-ISRIB compounds are predicted to increase eIF2B’s affinity for eIF2(αP) and thus extend the ISR, which may be of benefit in some contexts (for example as anti-viral agents). Given that, like ISRIB, a subset of such ligands may also accelerate the assembly of the eIF2B decamer (at least in vitro), their activity as ISR modulators may shed light on the relative role of these two known facets of ISRIB action in cells.

## Acknowledgements

We thank Steffen Preissler and Lisa Neidhardt (CIMR) for critiques of the manuscript, Alan Hinnebusch and Tom Dever (NIH/NICHD) for suggesting the experiment shown in figure 6C, Reiner Schulte, Chiara Cossetti and Gabriela Grondys-Kotarba (CIMR core flow cytometry facility) for their help and advice on cell sorting, and Akeel Abu Alard and Peter Fischer (U Nottingham) for the gift of ISRIB and FAM-ISRIB.

This work was supported by a Wellcome Trust Principal Research Fellowship to D.R. (Wellcome 200848/Z/16/Z), a Japan Society for the Promotion of Science (Grant-in-Aid for Scientific Research [B], [JP19H03172 to T.I.], Grants-in-Aid for Early-Career Scientists [JP18K14644 to K.K.]), Japan Agency for Medical Research and Development (Platform Project for Supporting Drug Discovery and Life Science Research [Basis for Supporting Innovative Drug Discovery and Life Science Research] [JP20am0101082]), and RIKEN (the Integrated life science research to challenge super aging society and the Pioneering Project “Dynamic Structural Biology”).

## Authors contribution

A.F.Z., K.K., T.I. and D.R. conceived the study, designed experiments, analysed data, prepared figures and tables, and co-wrote the manuscript. K.K., A.S., M.N., M.Y., M.S., A.F.Z, D.R. and L.A.P purified recombinant proteins. K.K., M.S. and T.I. conducted cryo-EM structural work. A.F.Z. and D.R. conducted GEF assays. D.R. conducted FP and BLI assays. L.A.P. assisted in the design and interpretation of the BLI experiments. A.F.Z. and H.H. conducted analytical gradient centrifugation of cell lysates. A.F.Z., A.C. and C.R. created and evaluated mutant cell lines. A.F.Z. and C.R. conducted CRISPR/Cas9-mediated depletion experiments in cells. All authors contributed to discussion and revision of the manuscript.

## Methods

### Protein preparation

Human eIF2B, wildtype, or ISR-defective δ^E310K^ or δ^L314Q^ mutant versions, were purified from a bacterial expression system, whereas human eIF2, wildtype or non-phosphorylatable α^S51A^ mutant, were purified from transfected FreeStyle 293-F cells as previously described (Kashiwagi et al., 2019). As the α and γ subunits of eIF2 in this study have C-terminal PA and FLAG-His_8_ tags, respectively, human eIF2 proteins were purified by a His-Accept column (Nacalai tesque), Anti PA tag Antibody Beads (Fujifilm Wako), and a HiTrap desalting column (GE Healthcare), and were dissolved in 20 mM HEPES-KOH buffer pH 7.5 containing 200 mM KCl, 1 mM MgCl_2_, 1 mM DTT, and 10%(^v^/v) glycerol. The C-terminally-AviTag and 6 x His tagged N-terminal lobe of human eIF2α (residues 1-187), was purified from bacteria (where it was biotinylated by the endogenous BirA) by Ni-NTA affinity chromatography.

Purified human eIF2 or the biotinylated N-terminal lobe of human eIF2α were phosphorylated in vitro in kinase buffer (20 mM HEPES-KOH pH 7.4, 150 mM KCl, 1 mM TCEP, 2 mM MgCl_2_, 1 mM ATP) using bacterial-expressed PERK kinase domain (immobilised on glutathione sepharose beads and removed from the reaction at conclusion by phase separation). Stoichiometric phosphorylation of the eIF2α subunit was confirmed on a Coomassie-stained PhosTag gel (see Figure 4, supplement 1A).

### Guanine nucleotide exchange activity

eIF2B guanine nucleotide exchange activity was measured as described previously (Sekine et al., 2015), with minor modifications. Briefly, purified decameric eIF2B (final concentration 2.5 - 40 nM), phosphorylated or unphosphorylated eIF2 (final concentration 0 - 1 μM), ISRIB (a gift of Peter Fischer, U, Nottingham) dissolved in DMSO (final concentration 0.25 - 1 μM) or DMSO carrier control (final concentration < 5%^v^/v) were pre-assembled in assay buffer (20 mM HEPES-KOH pH 7.4, 150 mM KCl, 2 mM MgCl_2_, 1 mM TCEP, 0.05 mg/mL bovine serum albumin, 0.01% Triton X-100, 1.5 mM GDP) and allowed to equilibrate for 10 minutes at room temperature in a low volume, “U” bottom, black 394 well plate (Corning, Cat #3667). At t = 0 purified eIF2(α^S51A^), preloaded with BODIPY-FL-GDP (Invitrogen) (as previously described, Sekine et al., 2015) was introduced at a final concentration of 125 nM and the fluorescence signal read kinetically in a Tecan F500 plate reader (Excitation wavelength: 485 nm, bandwidth 20 nm, Emission wavelength: 535 nm, bandwidth 25 nm). Where indicated, the data were fitted to a single-phase exponential decay function using GraphPad Prism v8.

### Cryo-EM analysis

Cryo-EM sample preparation and data collection were performed as previously described (Kashiwagi et al., 2019), but the ratio of eIF2B and eIF2(αP) was changed to 1:4, and they were diluted to 60 nM and 240 nM, respectively. The total number of collected images was 7,729.

The movie frames were aligned with MotionCor2 (Zheng et al., 2017) and the CTF parameters were estimated with Gctf (Zhang, 2016) in RELION-3.0 (Zivanov et al., 2018). To make the templates for automated particle picking with Gautomatch (http://www.mrc-lmb.cam.ac.uk/kzhang/), about 20,000 particles were semi-automatically picked with EMAN2 (Tang et al., 2007), and 2D averages were generated in RELION. Automatically picked 1,889,101 particles were extracted with rescaling to 2.94 Å/pix and 2D & 3D classification in RELION was performed. A low-pass filtered (40 Å) map calculated from the crystal structure of *Schizosaccharomyces pombe* eIF2B (PDB: 5B04) (Kashiwagi et al., 2016) was used as a reference map in 3D classification. After 3D classification steps, 365,487 particles in good classes were re-extracted without rescaling (1.47 Å/pix), and 3D refinement, Bayesian polishing, and CTF refinement were performed. These refined particles were applied to 3D classification again, and separated into classes in which one molecule of eIF2α (the α^P^1 complex, 208,728 particles, 3.8 Å), two molecules of eIF2α (the α^P^2 complex, 80,921 particles, 4.0 Å), or eIF2αγ at one side and eIF2α at the other side are resolved (the α^P^γ complex, 66,721 particles, 4.3 Å), respectively. In addition, the previous dataset for the eIF2BoeIF2(αP) complex (PDB: 6K72) (Kashiwagi et al., 2019) was also re-analysed. The 3D class not containing eIF2(αP) was selected re-extracted and refined as above (330,601 particles, 4.0 Å).

As a model, the cryo-EM structure of human eIF2B in complex with P-eIF2α at 3.0-Å resolution (PDB: 6O9Z) (Kenner et al, 2019) was used for the most part of eIF2B and the N-terminal domain of P-eIF2α. For the rest, the cryo-EM structure of human eIF2B in complex with eIF2(αP) at 4.6-Å resolution (PDB: 6K72) (Kashiwagi et al., 2019) was used. These structures were manually fitted into the maps. Map sharpening and model refinement were performed in PHENIX (Adams et al., 2010), and the models were further refined manually with Coot (Emsley et al., 2010). The refinement statistics of these structures are shown in Table S1.

### ISRIB binding to eIF2B

FAM-conjugated ISRIB (at 2.5 - 5 nM, final) (Zyryanova et al., 2018) was combined with purified eIF2B (6 - 150 nM) in presence or absence of phosphorylated or unphosphorylated eIF2 (final concentration 0 - 2.5 μM), the N-terminal lobe of phosphorylated eIF2α (final concentration 0 - 40 μM) or unlabelled ISRIB (0.5 - 1 μM) in assay buffer above, and allowed to equilibrate for 30 minutes at room temperature in a low volume, “U” bottom, black 394 well plate (Corning, Cat #3667). The fluorescence polarisation signal was read on a CLARIOstar microplate reader (BMG Labtech) with filter settings of 482 nm (excitation) and 530 nm (emission). Inhibition was fitted using the log(inhibitor) vs. response - variable slope (four parameters) function on GraphPad Prism v8.

Where indicated (Figure 4A) at t = 0 unlabelled ISRIB (1 μM final) or equal volume of DMSO carrier were introduced into samples containing pre-equilibrated FAM-ISRIB (a gift of Peter Fischer, U. Nottingham) and eIF2B (60 nM) and the fluorescence polarisation signal was read kinetically. The data were fitted to a single-phase exponential decay function using GraphPad Prism v8.

Where indicated (Figure 4B and figure 4, supplement 1A) at t = 0 PERK kinase (1 to 100 nM final concentration, of bacterially-expressed GST-PERK) was introduced into samples containing pre-equilibrated FAM-ISRIB (2.5 - 5 nM), eIF2B (60 - 83 nM), wildtype eIF2 or non-phosphorylatable eIF2(α^S51A^) (300 - 600 nM) in assay buffer supplemented with 1 mM ATP and the change in fluorescence polarisation was read kinetically.

Where indicated (Figure 4, supplement 1B), at t = 0 bacterially expressed lambda phosphatase (160 nM) or a pre-assembled complex of the trimeric eI F2(αP)-directed holophosphatase comprised of G-actin/PP1A catalytic subunit/PPP1R15A regulatory subunit (as described in Crespillo-Casado et al., 2018, final concentration, 100 nM G-actin, 100 nM PPP1R15A, 10 nM PP1A) was introduced in samples with pre-equilibrated FAM-ISRIB (2.5 nM), eIF2B (60 nM) and eIF2(αP) (300 nM) and the change in fluorescence polarisation was read kinetically.

### eIF2B binding to phosphorylated eIF2α

BLI experiments were conducted at 30°C on the FortéBio Octet RED96 System, at an orbital shake speed of 600 rpm, using Streptavidin (SA)-coated biosensors (Pall FortéBio) in 20 mM HEPES pH 7.4, 150 mM KCl, 2 mM MgCl_2_, 1 mM TCEP, 0.05 mg/mL bovine serum albumin and 0.01% Triton X-100. Biotinylated ligand [C-terminally-AviTag-His_6_ tagged N-terminal lobe of human eIF2α (residues 1-187 at a concentration of 150 nM)] was loaded to a binding signal of 1-2 nm, followed by baseline equilibration in buffer. Association reactions with analyte (wildtype or mutant eIF2B decamers or eIF2Bβδγε tetramers) prepared in the aforementioned buffer, or dissociation reactions in buffer, with ISRIB or an equal volume of DMSO were conducted with a reaction volume of 200 μL in 96-well microplates (greiner bio-one). In Figure 5C association reactions were conducted without ISRIB and the dissociation reactions (shown) were conducted with the indicated concentration of ISRIB, and equal final volumes of DMSO.

Data were analysed using Prism GraphPad V8, as indicated in the figure legends.

### Cell culture

CHO-K1-derived adherent cell lines were maintained in Nutrient Mixture F12 (N4888, Sigma), 10% Fetal Calf serum (FetalClone II, Thermo), 2 mM L-glutamine (G7513, Sigma Aldrich), and 1 x Penicillin/Streptomycin (P0781, Sigma) at 37°C with 5% CO_2_.

HeLa-derived adherent cell lines were maintained in DMEM (D6546, Sigma Aldrich) supplemented with 2 mM L-glutamine (G7513, Sigma Aldrich), 1 x Penicillin/ Streptomycin (P0781, Sigma), 1 x non-essential amino acids solution (M7145, Sigma), and 55 μM β-mercaptoethanol at 37°C with 5% CO_2_.

The list of new cell lines generated for the study is described in Table S2, the plasmids and primers used for transfection are in Tables S3 and S4, respectively.

### Measurement of the ISR in cultured cells

Generation of CHO-S21 and CHO-S21 *Eif2S1*^S51A^ cells containing a stably integrated ISR (CHOP::GFP) and UPR (Xbp1::Turquoise) responsive reporter was described previously (Sekine et al., 2016). Inhibition of histidyl-tRNA synthetase by histidinol in the parental CHO-S21 cells, but not in the *Eif2S1*^S51A^ mutant, activates the eIF2α kinase GCN2 that phosphorylates eIF2. eIF2(αP) inhibits its GEF eIF2B, initiating the ISR, and culminating in CHOP::GFP activation, which was detected by flow cytometry. In wild-type histdinol-treated cells the presence of ISRIB attenuates the response of the CHOP::GFP reporter, however, in eIF2(αS51A) mutant cells this effect can no longer be observed due to inability of histidinol to trigger the ISR response in those cells (also known as gcn-phenotype).

To observe the drugs effect in any of CHO cell lines, cells were split and seeded at confluency of 2 - 4 x 10^4^ cells/well on a 12-well plate. Two days later the medium was refreshed and cells were either treated with 0.5 mM L-histidinol (228830010, Acros Organics), or 200 nM ISRIB, or both for 18 - 24 hours. Immediately before flow cytometry analysis, cells were washed with ice-cold PBS, and collected in ice-cold PBS containing 4 mM EDTA pH 8.0. Fluorescent signal from single cells (10,000/ sample) was measured on LSRFortessa cell analyser.

The populations of cells were further analysed on FlowJo software where the median for each Gaussian distribution was defined. For the samples containing bimodal distribution two medians were defined. To assess the ISR folds increase in each transfected sample the median of the “ISR-on” CHOP::GFP signal (right distribution) was divided by the median of the “ISR-off” CHOP::GFP signal (left distribution). In the case of a unimodal distribution the median of a given population was divided on itself. The means of three repeats with standard deviations and P values were obtained using Prism software.

### eIF2B and eIF2 subunits depletion

CHO-S21 *Eif2S1*^S51A^ cells were split and seeded at density of 5 x 10^4^ cells/well on a 12-well plate. The next day cells of about 20-30% confluence were pre-treated for 60 minutes with either 1 μM of ISRIB in DMSO or the equivalent amount of 100% DMSO and then transfected with 1 μg of CRISPR/ Cas9 plasmid either without (control) of with sgRNA (see Table S3 for plasmids and Table S4 for primers) using Lipofectamine LTX with Plus Reagent (A12621, Thermofisher) according to the manufacturer’s protocol. Medium supplemented with either 1 μM ISRIB or the equivalent amount of 100% DMSO was refreshed every 24 hours thereafter. On the day of harvest (48, 72 and 96 hours post transfection) cells were washed twice with ice-cold PBS, harvested in 0.5 mL of ice-cold PBS supplemented with 4 mM EDTA and immediately analysed on LSRFortessa cell analyser. CHOP::GFP (excitation 488 nm/ emission 530 ± 30 nm) fluorescence signal from single cells (20,000/ sample) was measured. The populations of cells were further analysed on FlowJo software as described above.

### Introduction of ISR resistant mutations into cultured cells

Assessing the importance of counterparts to *S. cerevisiae* eIF2Bδ residues GCD2^E377^ and GCD2^L381^ (known ΔISR/ΔGCN yeast mutants, Pavitt et al., 1997’, E312 and L316 in the hamster genome, and E310 and L314 in the human) to the ability of eIF2B to respond to eIF2(αP) and initiate an ISR in vivo, was carried out by targeting the *Eif2b4* locus of CHOP::GFP carrying CHO-S21 cells (described above) with a CRISPR/Cas9 guide (GAAGATTGTGCTTGCAGCTC**AGG**, PAM sequence in bold) and providing an ssODN repair template randomised at codon L316 and either carrying the wildtype sequence at E312 (oligo #2213, eIF2B4_ ssODN_L316X) or an additional E312K mutation (oligo #2214, eIF2B4_ ssODN_E312K_L316X) (Table S4). The transduced cells were selected for ISR resistance based on defective CHOP::GFP induction in response to histidinol. Single clones were sequenced, two of which 12H6 (genotype *Eif2b4^L316N^*) and 22H2 (genotype *Eif2b4^E312K; L316V^*) were selected for further study (Table S2 and Figure 1C).

The effect of the mutations on translational control in response to stress was assessed by measuring the incorporation of puromycin into newly synthesized proteins by immunoblotting lysates of untreated and thapsigargin (Sigma, T9033) (200 nM, 45’)-treated cells that had been exposed to 10 μg/mL puromycin (Sigma, P8833) 10 minutes before lysis. Immunoblot detection was conducted using primary antibodies for puromycinylated protein (Schmidt et al., 2009), phospho-eIF2α-Ser51 (Epitomics), or total eIF2α (Scorsone et al., 1987), and IR800 or IR680 conjugated secondary antisera followed by scanning on a Licor Odyssey scanner. The extent of the ISR defect was benchmarked against CHO-S21 cells with an *Eif2S1^S51A^* knockin mutation (Sekine et al., 2016)

### Glycerol gradient fractionation of cell lysates

CHO-S7 [with a 3XFLAG tag knocked into their *Eif2b3* locus (*Eif2b3*^3xFLAG in/+^ Sekine et al., 2016)] and HeLa [with a 3XFLAG tag knocked into their *EIF2B2* locus (*EIF2B2*^3xFLAG in/in^ Sekine et al., 2015)] cells (9 x 10^7^ cells/ sample) were harvested, lysed in 250-500 μL of lysis buffer (50 mM HEPES-KOH pH 7.5, 150 mM NaCl, 1% (v/v) Triton, 5% (v/v) Glycerol, 1mM DTT, 2 mM PMSF, 8 μg/ml aprotinin, 4 μg/mL pepstatin) either with 250 nM ISRIB (in DMSO) or equivalent amount of 100% DMSO, and cleared supernatant was applied on 5 mL of 10 - 40% (v/v) glycerol gradient prepared in lysis buffer (without triton) with respective amounts of glycerol using SG15 Hoefer Gradient Maker and centrifuged using SW50 (Beckman Coulter) rotor at 45,000 rpm for either 13 hours or for 14 hours 48 minutes at 4°C. After the centrifugation gradients were manually fractionated into 16 fractions of 325 μL, and 30 μL of each fraction was taken for western blot analysis. Fractions were run on 10% SDS-PAGE gel, transferred onto PVDF membrane, incubated for 2 hours at RT with primary monoclonal mouse anti-FLAG M2 antibody (F1804, Sigma Aldrich) to track migration of 3 x FLAG-tagged eIF2B complex, followed by incubation for 45 min at RT with secondary goat anti-mouse-HRP antibodies according to the manufacturer’s protocol. Membranes were developed with enhanced chemiluminescence kit following the manufacturer’s procedure, and scanned on CheminDoc (Bio-Rad). Image analysis was done using ImageJ software.

## SUPPLEMENTAL MATERIALS

**Table S1.**
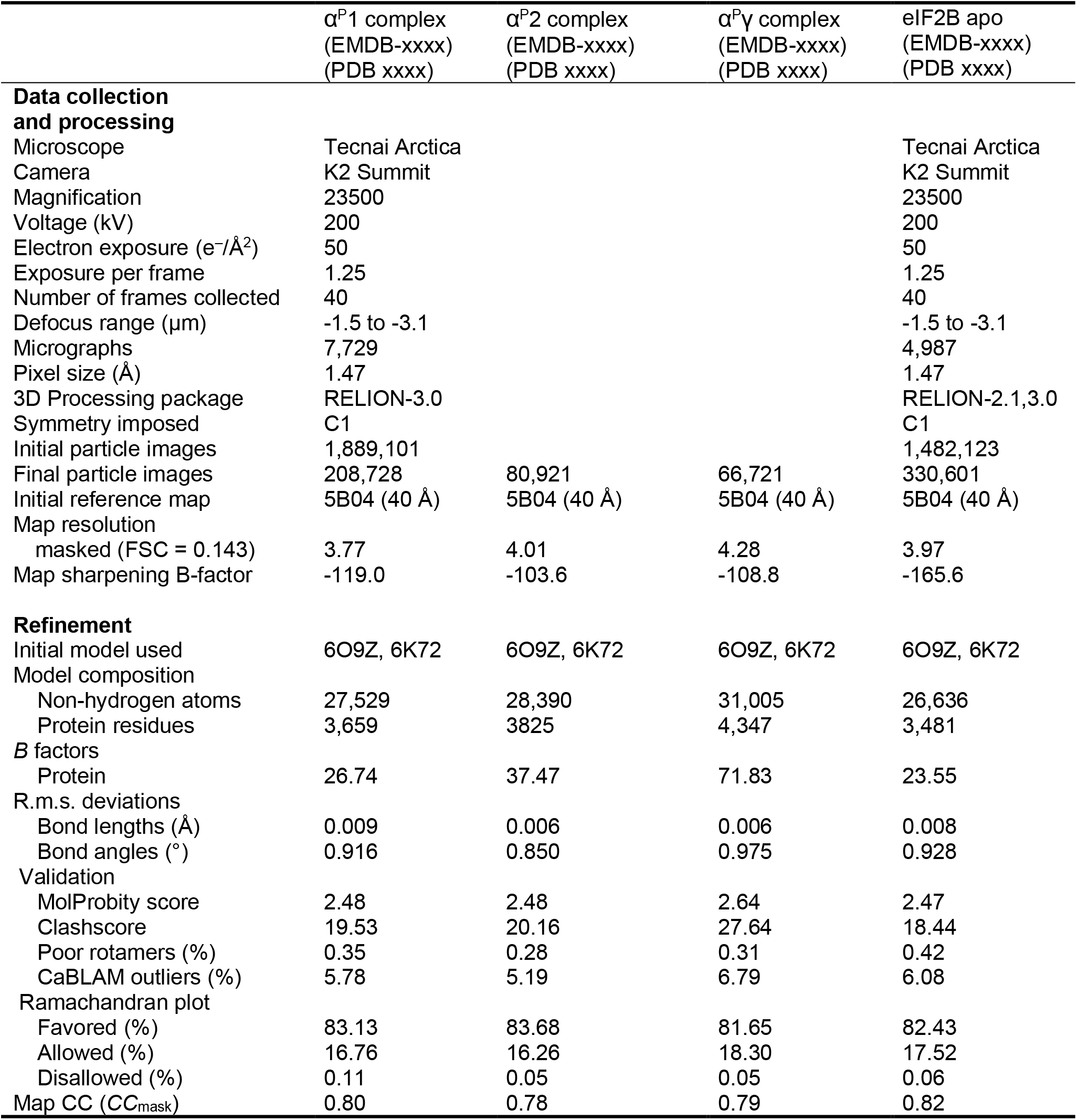
Cryo-EM data collection and image processing.

**Table S2.** List of cell lines.

| Gene | Exon | Cells | Clone name | Description | Mutagenized region (numbers indicate amino acid position at which mutagenesis occurred) |
| --- | --- | --- | --- | --- | --- |
| NA | NA | CHO-S21 | NA | dual reporter [CHOP::GFP; Xbp1::Turquoise] parental cell line from Sekine et al. 2016 | NA |
| *Eif2S1* | 2 | CHO S51A | NA | dual reporter ISR-insensitive (gcn-) eIF2αS51A from Sekine et al. 2016 | 51_ <u>A</u>RRRIRSI |
| *Eif2b4* | 10 | CHO-S21 | 12H6 | eIF2Bδ(L316N), ISR-insensitive | 316_ <u>N</u>AAQAISRF |
| *Eif2b4* | 10 | CHO-S21 | 22H2 | eIF2Bδ(E312K; L316V), ISR-insensitive | 312_ <u>K</u>KIV_316_ <u>V</u>AAQA |

**Table S3.** List of plasmids.

| ID | Plasmid name | Description | Primers used to generate plasmid | Figures |
| --- | --- | --- | --- | --- |
| UK2320 | CHO_EIF2B4_EXON10_g3_pSpCas9(BB)-2A-Puro | CRISPR/Cas9 with puromycin selection targeting hamster *Eif2b4* (eIF2B delta) gene | Oligo 2209 & 2210 | 1C |
| UK2733 | heIF2a_2-187_WT_AviTag_H6_pET-30a(+) | wildtype NTD human eIF2α_1-187 with AviTag and 6x histidines in bacterial expression vector | NA | 3B |
| UK1610 | pSpCas9(BB)-2A-mCherry_V2 | CRISPR/Cas9 empty vector with mCherry selection | NA | 5A |
| UK2536 | cgelF2B2_g2_pSpCas9(BB)-2A-mCherry | CRISPR/Cas9 with mCherry selection targeting hamster *Eif2b2* (eIF2B beta) gene (guide 1) | Oligo 2520 & 2521 | 5A |
| UK2537 | cgelF2B2_g3_pSpCas9(BB)-2A-mCherry | CRISPR/Cas9 with mCherry selection targeting hamster *Eif2b2* (eIF2B beta) gene (guide 2) | Oligo 2522 & 2523 | 5A |
| UK2538 | cgelF2B4_g1_pSpCas9(BB)-2A-mCherry | CRISPR/Cas9 with mCherry selection targeting hamster *Eif2b4* (eIF2B delta) gene (guide 1) | Oligo 2524 & 2525 | 5A |
| UK2539 | cgelF2B4_g3_pSpCas9(BB)-2A-mCherry | CRISPR/Cas9 with mCherry selection targeting hamster *Eif2b4* (eIF2B delta) gene (guide 2) | Oligo 2526 & 2527 | 5A |
| UK2547 | cgelF2B5_g1_pSpCas9(BB)-2A-mCherry | CRISPR/Cas9 with mCherry selection targeting hamster *Eif2b5* (eIF2B epsilon) gene | Oligo 2543 & 2544 | 5A |
| UK1367 | pSpCas9(BB)-2A-Puro | CRISPR/Cas9 empty vector with puromycin selection | NA | 5C |
| UK1505 | CHO_Eif2s1_guideA_pSpCas9(BB)-2A-Puro | CRISPR/Cas9 with puromycin selection targeting hamster *Eif2s1* (eIF2 alpha) gene (guide A) | Oligo 1015 & 1018 | 5C |
| UK1506 | CHO_Eif2s1_guideB_pSpCas9(BB)-2A-Puro | CRISPR/Cas9 with puromycin selection targeting hamster *Eif2s1* (eIF2 alpha) gene (guide B) | Oligo 1016 & 1019 | 5C |

**Table S4.**
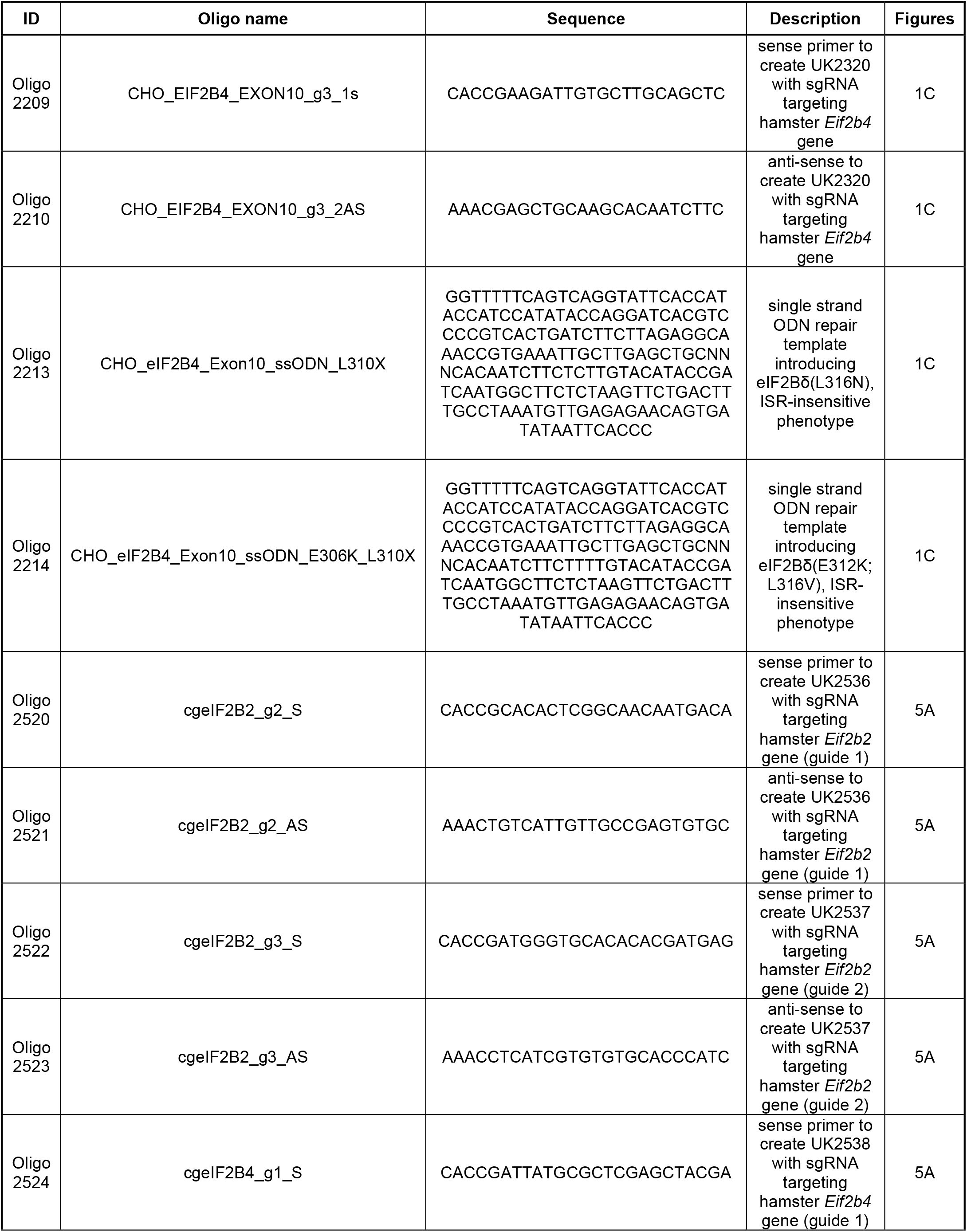
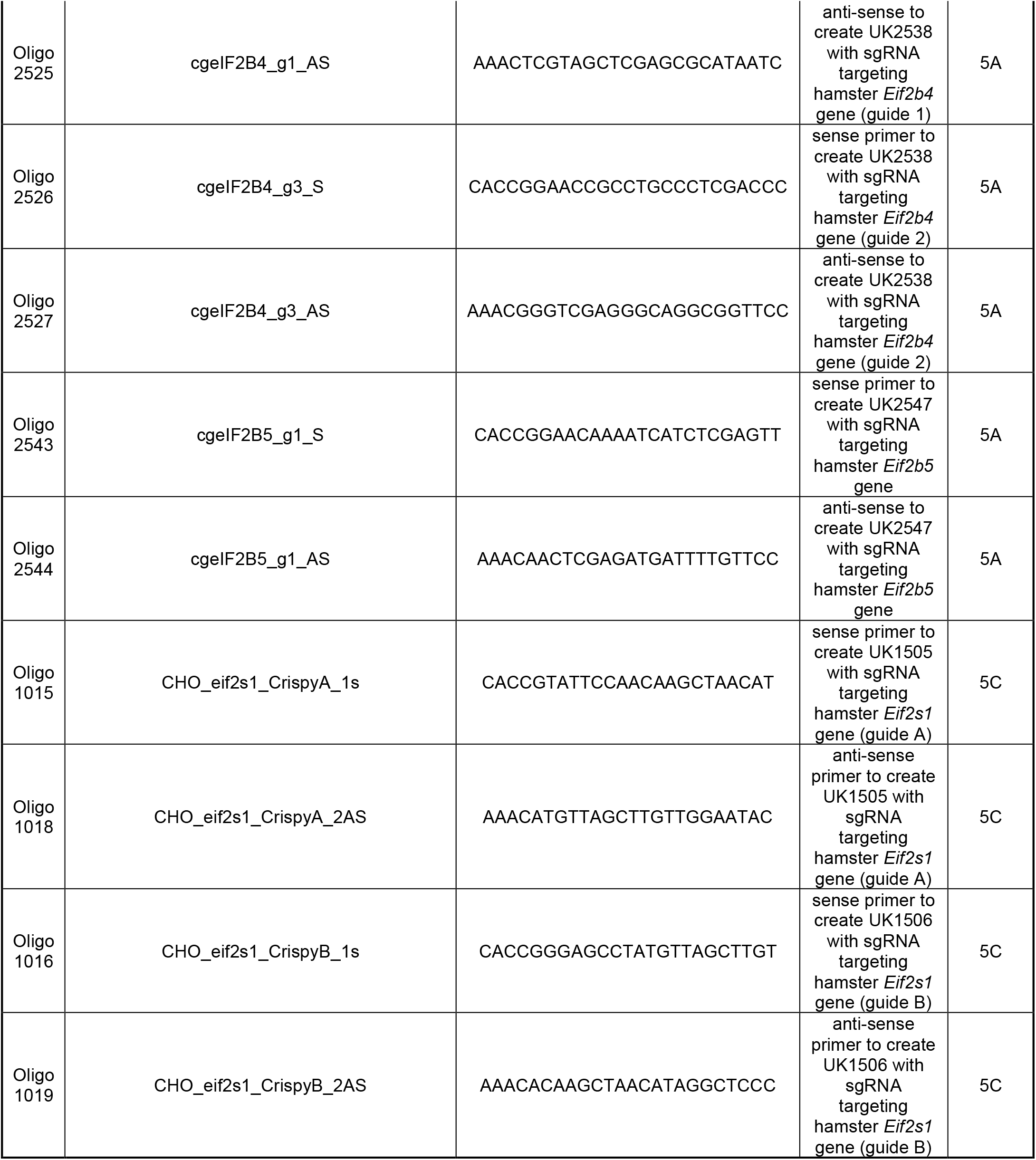
List of primers.

---

**Figure 2 supplement 1.**
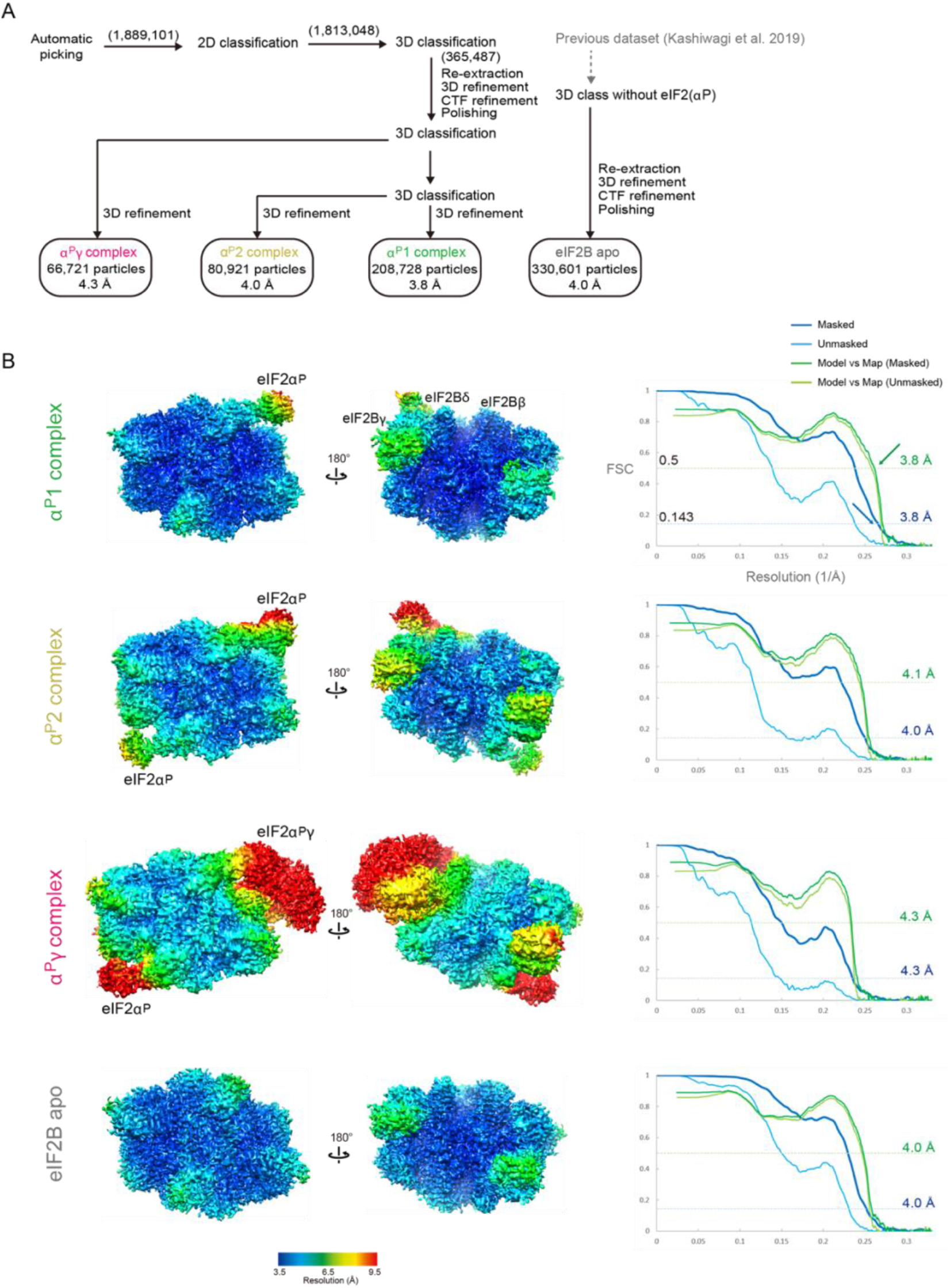
Cryo-EM data processing. A) Workflow of image processing. Total particle numbers at each stage are show in parentheses. B) Local resolution maps and Fourier shell correlation (FSC) curves of the cryo-EM maps. The FSC curves for masked (blue), unmasked (cyan) map, and the curves for model and map correlation (masked: green, unmasked: yellow green) are shown. The resolutions at which FSC for masked map drops below 0.143 and model map correlation drops below 0.5 are shown.

**Figure 2 supplement 2.**
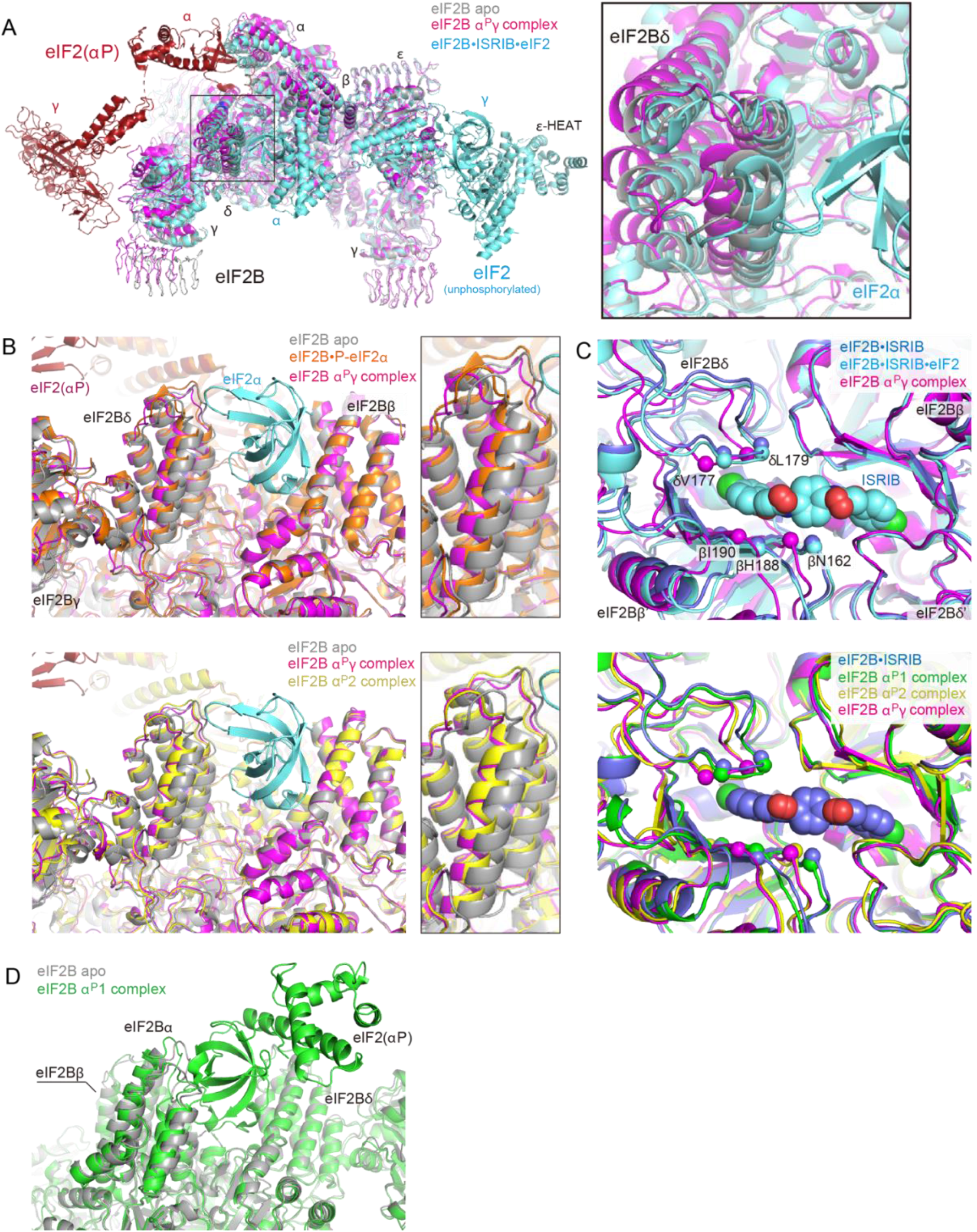
Changes to the catalytically productive interface and the ISRIB-binding pocket of eIF2B induced by eIF2(αP) binding. A) Overlay of the eIF2B apo structure (grey), eIF2B in complex with two eIF2(αP) trimers (the α^P^γ complex; eIF2B in pink and eIF2(αP) in red), and eIF2B•ISRIB•eIF2 complex (cyan, PDB: 6O85) (same view as Fig. 2A). Structures are aligned by their eIF2Bβ subunits. Right panel: Close-up view of the movement of eIF2Bδ by the binding of the unphosphorylated (cyan) or phosphorylated eIF2 (pink). B) Views of the catalytically productive interface in apo eIF2B compared to complexes with the isolated phosphorylated eIF2α subunit (P-eIF2α) or eIF2(αP) trimer in which the eIF2βγ lobe is not (α^P^2) or is resolved (α^P^γ). Note that the δ subunit of the eIF2BoP-eIF2α complex structure (orange in A, PDB: 6O9Z) resides in a position intermediate between eIF2B apo structure (grey) and the α^P^γ complex structure (pink). The α^P^2 complex (yellow in B) induces a similar displacement as the α^P^γ complex. The N-terminal S1 domain of eIF2α in the productive eIF2B•ISRIB•eIF2 complex is shown as reference (cyan, PDB: 6O85). C) Views of the ISRIB-binding pocket. Note the similarity in disposition of key residues involved in ISRIB action in the eIF2BoISRIB (blue, PDB: 6CAJ) and eIF2B•ISRIB•eIF2 complex (cyan, PDB: 6O85), and the deformation of the pocket brought about by binding two phosphorylated eIF2 trimers (the α^P^γ complex in pink and the α^P^2 complex in yellow), and the intermediate conformation imposed by binding of one phosphorylated eIF2 trimer (the α^P^1 complex in green). Key residues lining the pocket, including eIF2B δL179 and βH188 that are known to affect the binding or action of ISRIB, are highlighted as spheres. Structures are aligned to the δ’ subunit of the eIF2BoISRIB complex. D) Movement of the eIF2Bα subunit in the α^P^1 complex (green) relative to eIF2Bα subunit in eIF2B apo structure (grey). Structures are aligned by their eIF2Bβ subunits.

**Figure 4 supplement 1.**
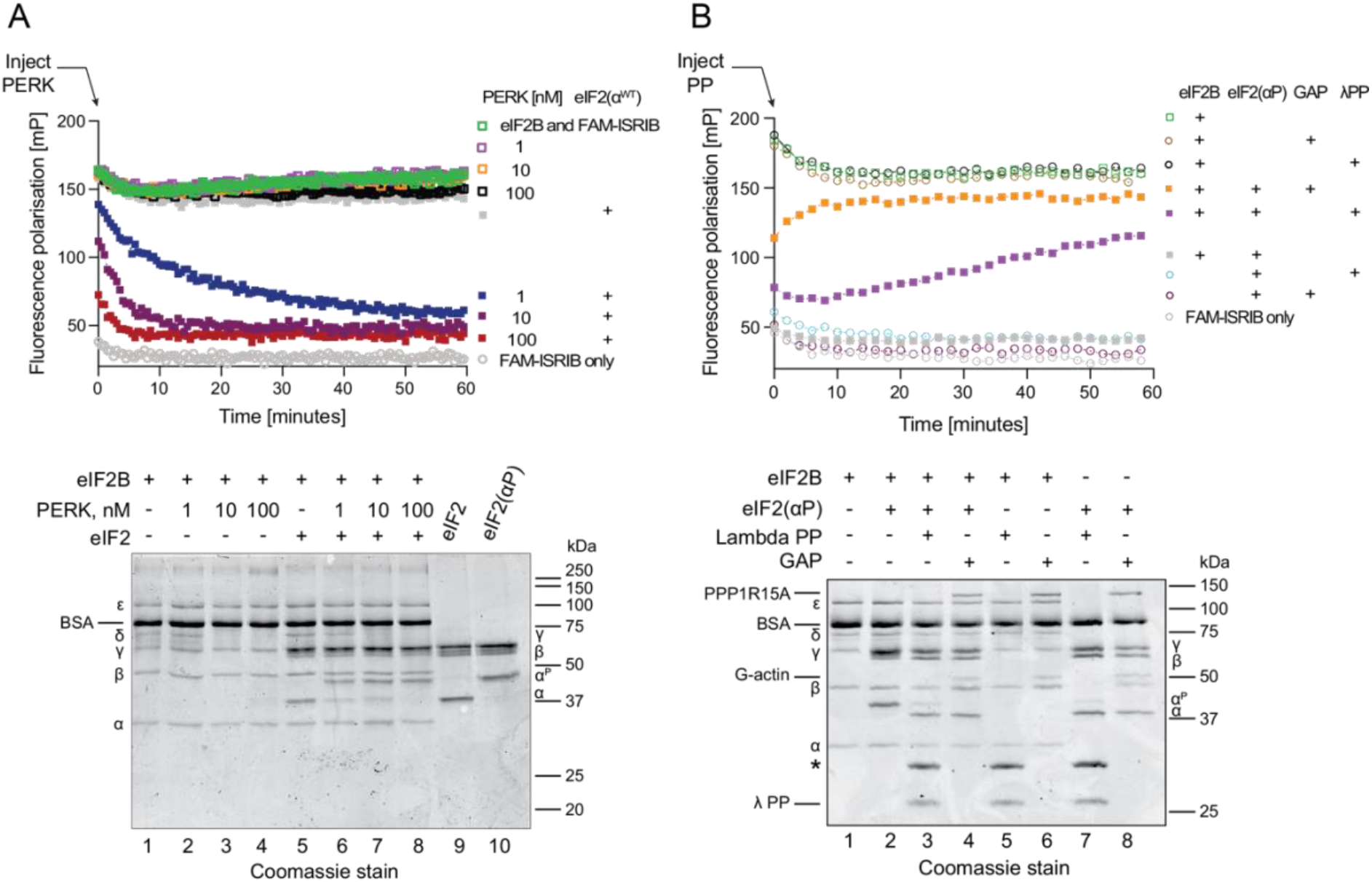
eIF2(αP)-mediated inhibition of FAM-ISRIB binding to eIF2B is captured kinetically and is reversible by dephosphorylation. A) Upper left panel: plot of time-dependent change in fluorescence polarisation of FAM-ISRIB bound to wildtype eIF2B in presence or absence of unphosphorylated eIF2. Where indicated, at t = 0 the eIF2α kinase PERK was introduced at varying concentrations to promote a pool of eIF2(αP). Shown is a representative experiment (one of three). Lower left panel: Coomassie stained PhosTag SDS-PAGE of the samples analysed in the experiment above. Migration of the eIF2 subunits, including phosphorylated and nonphosphorylated eIF2α, are indicated on the right. Pure samples of unphosphorylated and phosphorylated eIF2 are provided as references. The prominent band at ~70 kDa present in all lanes is bovine serum albumin (BSA), utilised as a stabiliser in all reactions (it obscures the GST-PERK signal, where applicable). Migration of eIF2B subunits is indicated on the left. B) Upper right panel: Plot of time-dependent change in fluorescence polarisation of FAM-ISRIB bound to wildtype eIF2B in presence or absence of phosphorylated eIF2. Where indicated, at t = 0 a specific eIF2(αP)-directed holophosphatase consisting of G-actin/PP1A/PPP1R15A (GAP) or the non-specific lambda phosphatase (λP) was introduced to convert phosphorylated eIF2 to eIF2. Shown is a representative experiment (one of two). Lower right panel: Coomassie stained PhosTag SDS-PAGE gel of the samples analysed in the experiment above. The eIF2 subunit, including phosphorylated and unphosphorylated eIF2α, are indicated on the right, eIF2B subunits and species arising from the phosphatase-treated samples are indicated on the left (the catalytic subunit PP1A is not visible on this gel; the asterisk marks an unidentified contaminant of the λP samples).

